# Mutagenic bypass of uracil-derived abasic sites underlies the SBS17 mutational signature

**DOI:** 10.64898/2026.08.21.746158

**Authors:** Stefania Di Ciccio, Joeri van Strien, Yang Jiang, Ammal Abbasi, Linda Bakker, Naomi Weertman, Laurynas Leiteris, Jeroen Willems, Ludmil B. Alexandrov, Juan I. Garaycoechea

## Abstract

Understanding how mutations arise is central to predicting and preventing cancer, yet most of the mutational signatures found in cancer genomes still have no known mechanistic cause. Among these, single-base-substitution signature 17 (SBS17) is the dominant mutational process in gastric and oesophageal adenocarcinoma. SBS17 has been linked to oxidative damage and to the chemotherapeutic agent 5-fluorouracil (5-FU), but the molecular events that generate it are unknown. Here we demonstrate that SBS17 causes driver mutations in gastrointestinal cancer. We then combine genetics in cell lines and organoids with whole-genome sequencing to define the mechanistic aetiology of SBS17 mutagenesis. We disprove the prevailing oxidative damage hypothesis and instead show that SBS17 arises through the misincorporation of dUTP during DNA replication, followed by uracil excision by the glycosylase UNG and mutagenic bypass of the resulting abasic (AP) sites by the translesion synthesis machinery. Importantly, we reveal that the same mechanism underlies both spontaneous and chemotherapy-induced mutations. Together, our results resolve the origin of gastric mutagenesis, opening therapeutic avenues for curbing ongoing mutagenesis, tumour evolution and drug resistance in gastrointestinal cancers.

## Main

Throughout life, every cell accumulates somatic mutations that arise from a continual interplay between DNA damage and the repair pathways that oppose it. Most of these mutations are functionally inert, but the occasional alteration of a cancer gene can initiate and drive tumour development^1,2^. Because each mutagenic process alters the genome in a chemically specific way, it leaves behind a characteristic pattern known as a mutational signature^3,4^. Mutational signatures therefore record the cumulative imprint of the DNA damage and repair processes that operate over a cell’s lifetime. Among them, single-base-substitution signature 17 (SBS17) is one of the most distinctive (**Fig. 1a**). The pattern now recognized as SBS17 was first reported in oesophageal adenocarcinoma^5^ and was subsequently found to be prevalent in oesophageal and gastric adenocarcinomas^4^. Later analyses resolved SBS17 into two components: SBS17a comprising T>A and T>C substitutions in a 5’C<u>T</u>N’3 context, and SBS17b comprising T>G substitutions in a 5’N<u>T</u>T’3 sequence context (**Fig. 1a**)^4,6^. In oesophageal and gastric adenocarcinomas, SBS17 can account for a substantial fraction of the mutational burden and is associated with *TP53* loss, chromosomal instability and disease progression^7,8^. Yet it also appears sporadically, and usually at low levels, across a range of other tumour types^4^. Strikingly, the signature is largely absent from normal somatic tissues^9^; its earliest appearance is in Barrett’s oesophagus (the metaplastic precursor of oesophageal adenocarcinoma), where it emerges in chromosomally unstable cells but not in chromosomally stable or non-diseased control cells, marking SBS17 as an early event in the neoplastic progression^7,8^. The molecular basis of SBS17 mutagenesis remains unknown, and defining it is a prerequisite to understanding why this process predominates in the upper gastrointestinal tract.

**Fig. 1.**
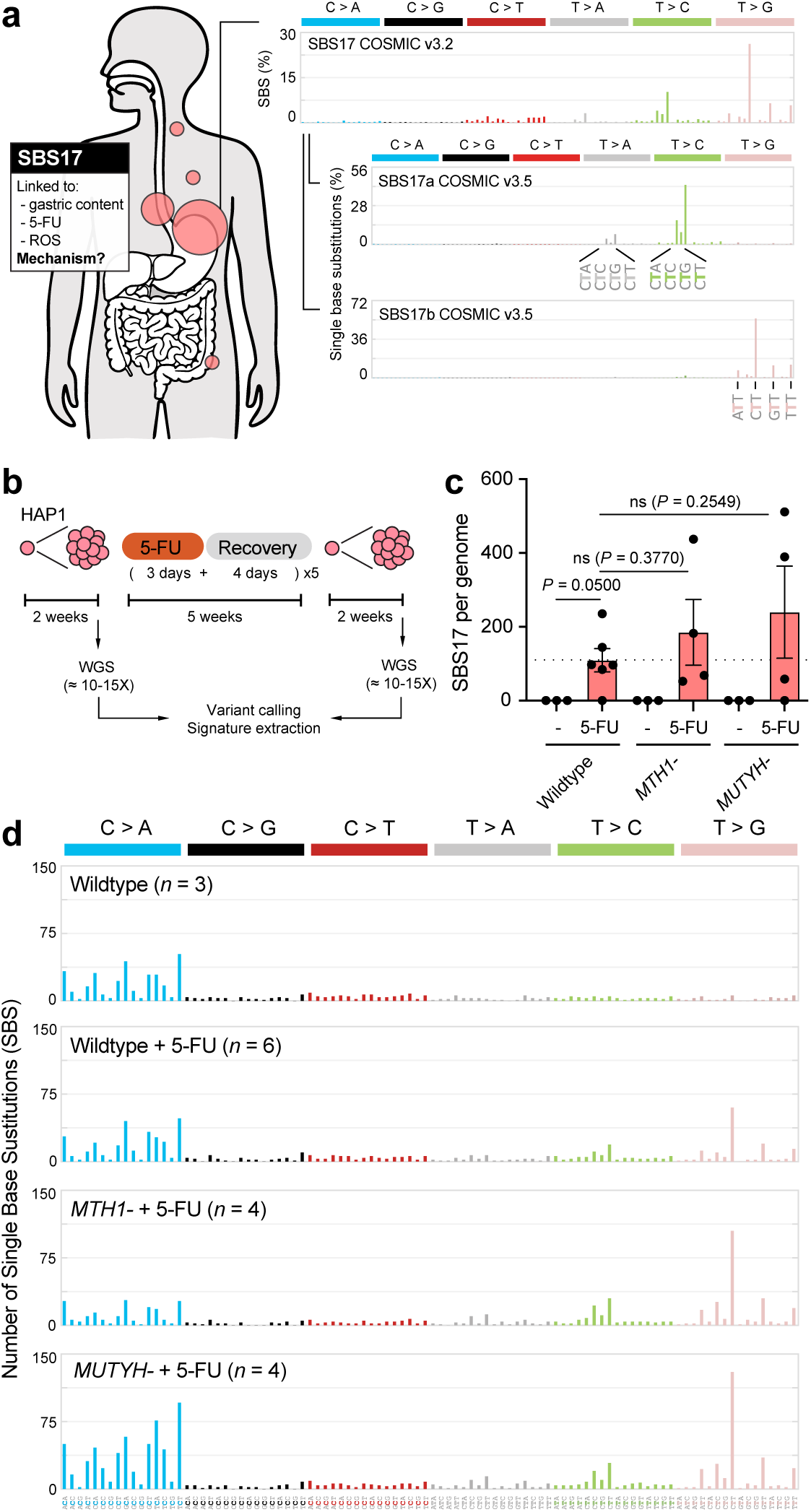
SBS17 is not driven by oxidation of the dGTP pool. **a)** Single Base Substitution signature 17 (SBS17) and its components SBS17a and SBS17b. The 96-classes of single-base substitutions (SBSs) consider the six mutation types but also the bases immediately 5’ and 3’ of the mutated base. SBS17 is predominantly found in oesophageal and gastric carcinoma. SBS17 has been linked to gastric content, reactive oxygen species (ROS) and the chemotherapeutic 5-fluorouracil (5-FU) but its molecular mechanism is currently unknown. **b)** *In vitro* experimental set up to induce SBS17 mutations with 5 cycles of 5-FU treatment. **c)** Burden of SBS17 mutations per (haploid) genome (*P* calculated by two-tailed unpaired *t* test, data shown as mean and s.e.m., *n* = 3-6). **d)** 96-classes of SBS considering the six mutation types but also the bases immediately 5’ and 3’ of the mutated base. Each graph represents the average mutation pattern for *n* genomes, where *n* is indicated in each panel.

A prevailing hypothesis attributes SBS17 to oxidative stress^10–12^. The upper gastrointestinal epithelium is chronically exposed to inflammation, reflux and bile acids, conditions that can promote the generation of reactive oxygen species (ROS)^13^. Whereas oxidation of guanine on DNA predominantly generates C>A mutations, oxidation of free dGTP produces 8-oxo-dGTP, which can be misincorporated opposite template adenine during replication to produce T>G transversions, as demonstrated in *E. coli*^14,15^. However, direct experimental evidence for this model is lacking, and 8-oxo-dGTP misincorporation produces only T>G substitutions, accounting at most for SBS17b and unable to generate the T>A and T>C mutations that define SBS17a.

SBS17 mutations have also been linked to the chemotherapeutic 5-fluorouracil (5-FU), a cornerstone of standard-of-care treatment for gastrointestinal cancers^16,17^. A drug with multiple cytotoxic effects, 5-FU perturbs nucleotide metabolism and itself incorporates into nucleic acids. Exposure of normal intestinal organoids to 5-FU *in vitro*, and analysis of 5-FU–treated colorectal and breast tumours *in vivo*, both yield SBS17 (refs.18,19). It has been suggested that 5-FU might likewise drive the T>G mutations of SBS17b by inducing ROS^10–12,20,21^. However, this model does not explain how the canonical antimetabolite activity of 5-FU relates to SBS17 mutagenesis and again leaves the T>A and T>C mutations of SBS17a unexplained. That an exogenous drug and an endogenous process should nonetheless converge on the same highly characteristic pattern raises the possibility that they share a common molecular intermediate.

Despite these clues, the mechanism that generates SBS17 remains undefined. Resolving this is central to understanding why SBS17 dominates oesophageal and gastric cancer. Here we combine CRISPR-engineered cell lines and organoid models with whole genome sequencing to uncover the molecular events leading to the generation of SBS17 mutations.

### SBS17 contributes to driver mutations in gastrointestinal cancers

We first examined the prevalence and contribution of SBS17 across 8,795 primary and 5,787 metastatic tumours profiled by whole-genome sequencing. SBS17 was detected across multiple tumour types but was particularly prevalent in oesophageal adenocarcinoma, occurring in approximately 87% of primary and 81% of metastatic tumours, and in stomach cancer, where it was present in approximately 67% of primary and 81% of metastatic tumours (**Extended Data Fig. 1a**). The largest change in prevalence was observed in colorectal cancer, increasing significantly from approximately 18% in primary tumours to 50% in metastatic disease. Breast cancer also showed a similar increase from approximately 1% to 10%, and significant primary-to-metastatic prevalence changes were also observed in several other tumour types, including pancreas, bladder, lung, head and neck, prostate and ovary. When present, SBS17 constituted a substantial fraction of the mutational burden, with median contributions of approximately 29% of substitutions in primary oesophageal adenocarcinoma, 23% in stomach cancer and 10% in colorectal cancer. Among SBS17-positive tumours, absolute SBS17 burden increased significantly from primary to metastatic oesophageal adenocarcinoma (1.53-fold, *q* = 6.7 × 10^−5^), colorectal cancer (1.59-fold, *q* = 0.0012) and breast cancer (3.9-fold, *q* = 9.7 × 10^−4^), with no significant change in stomach cancer (**Extended Data Fig. 1b**).

Consistent with experimental evidence that 5-FU induces SBS17 ^19^, documented 5-FU or capecitabine exposure was associated with a significantly higher prevalence of both SBS17a and SBS17b in metastatic colorectal and breast cancers (all q < 0.01). Notably, fluoropyrimidine exposure was associated more strongly with the presence of SBS17 than with its burden; among tumours already positive for SBS17 signatures, there were no significant differences in burden between exposed and unexposed tumours. This association may contribute to the marked primary-to-metastatic increases in SBS17 prevalence observed in colorectal and breast cancers, whereas SBS17 prevalence was already high among unexposed metastatic oesophageal and stomach cancers (**Extended Data Fig. 1c**).

We next asked whether SBS17 contributes to cancer-driver mutations. In Barrett’s oesophagus, SBS17 burden was 1.87-fold higher in progressors than non-progressors (*P* = 0.094), and SBS17-attributed drivers were detected in 7 of 40 (17.5%) progressors compared with 2 of 40 (5%) non-progressors, with neither comparison reaching statistical significance (**Extended Data Fig. 1d**). SBS17-attributed drivers were also detected in oesophageal adenocarcinoma, stomach and colorectal cancers, with a significant increase from primary to metastatic colorectal cancer (*q* = 0.009; **Extended Data Fig. 1e**). At the gene level, the strongest SBS17 attribution was observed for *CD5L* in Barrett’s progressors and *ERBB2* in metastatic oesophageal adenocarcinoma (**Extended Data Fig. 1f,g**). Notably, the recurrent ERBB2 S310F and V777L mutations were not attributed to SBS17, whereas SBS17 attribution was concentrated among non-recurrent *ERBB2* driver mutations.

Given the prevalence and contribution of SBS17 to driver mutations in gastrointestinal cancers, together with evidence that 5-FU exposure can generate the signature in some tumour contexts, we next sought to define its molecular mechanism.

### SBS17 mutations are not caused by oxidation of the dGTP pool

To dissect the molecular mechanism underlying SBS17 mutations, we established a cellular system to induce them in the human haploid cell line HAP1, using five cycles of 5-FU treatment as previously reported for organoids^18^. A single-cell–derived clone was split into four replicate cultures, each subjected to cycles of 3 days of 5-FU 10 μM followed by 4 days of recovery. After five treatment cycles, we expanded single haploid cells into clonal cultures and whole-genome sequenced them to 10–15X coverage (**Fig. 1b**). Following sequencing, variant calling and signature assignment, we found 5-FU treatment generated approximately 100 SBS17 mutations (defined as SBS17a + SBS17b) per haploid wildtype genome (**Fig. 1c,d**).

We first investigated the prevailing model that SBS17 arises through oxidation of the free dGTP pool. 5-FU has been shown to induce ROS^20,21^ and in *E. coli*, oxidation of the dGTP pool is a well-established mutagenic process^14,15^: 8-oxo-dGTP is incorporated opposite template adenine, giving rise to T>G mutations (**Extended Data Fig. 2a**). This is normally suppressed by the sanitising enzyme MutT, which hydrolyses 8-oxo-dGTP to 8-oxo-dGMP to prevent its incorporation; consequently, *mutT*-null *E. coli* displays a T>G hypermutator phenotype^14,15^. Once 8-oxo-dGTP is incorporated, the glycosylase MutY further promotes T>G mutations by excising the template adenine paired with the newly incorporated 8-oxo-G. To test the ROS hypothesis in mammalian cells we knocked out the mammalian homologues of MutT (*MTH1*) and MutY (*MUTYH*) (**Extended Data Fig. 2b,c**) and repeated the mutation-accumulation experiment. Neither perturbation significantly altered SBS17 burden (**Fig. 1c**), while both SBS17a and SBS17b remained evident in the resulting spectra (**Fig. 1d**).

In a subsequent experiment, we addressed potential redundancy. In mammals, MTH1 is the orthologue of MutT but additional homologues from the NUDIX family (MTH2, MTH3 and NUDT5) may have functional redundancy^22–26^. Additionally, in *E. coli* the glycosylase MutM (*OGG1*) acts downstream of MutY (*MUTYH*) to promote T>G mutations^15^. To test for redundancy between these factors, we generated a quadruple *MTH1^−^MTH2^−^MTH3^−^NUDT5^−^*and a double *MUTYH^−^OGG1^−^* knockout (**Extended Data Fig. 2b,c**), and repeated the mutation accumulation experiment. Neither genotype produced a statistically significant change in SBS17 induction by 5-FU, although the limited replicate numbers cannot exclude more modest effects (**Extended Data Fig. 3a-c**). Other perturbations nevertheless produced the expected phenotype: untreated *MUTYH^−^* cells showed an increased in C>A mutations resembling SBS36 (refs.^27,28^), consistent with oxidation of guanine on DNA, and C>A mutations were further exacerbated by loss of *OGG1* (**Extended Data Fig. 3d**). Increasing oxygen tension from 5% to 21% likewise significantly increased C>A mutagenesis, confirming increased oxidative damage on DNA, yet produced no significant increase in T>G mutations, including in *MTH1^−^MTH2^−^MTH3^−^NUDT5^−^* cells, arguing against 8-oxo-dGTP incorporation being a significant source of mutation in mammalian cells (**Extended Data Fig. 3e**). An independent experiment additionally generated a C>A-rich signature of uncertain origin; because this signature was experiment-specific, the datasets were analysed separately (**Extended Data Fig. 3b,f**). Taken together, these results argue against 8-oxo-dGTP being the source of T>G mutations following 5-FU treatment in HAP1 cells.

### SBS17 mutations are induced by abnormal dUTP pools

Having excluded oxidation of the dGTP pool, we next considered other effects of 5-FU, which can enter three major pathways (**Fig. 2a**). First, the majority of 5-FU is converted to the ribonucleotide 5F-UMP and ultimately 5F-UTP, which is incorporated into RNA and considered a principal source of its cytotoxicity^16,29^. Second, a smaller fraction is metabolised to the 2′-deoxynucleotide 5F-dUMP, a potent inhibitor of thymidylate synthase (TYMS), which catalyses the conversion of dUMP to dTMP; TYMS inhibition imbalances the dUTP/dTTP pool and depletes the dTTP needed for DNA synthesis. Finally, 5F-dUMP can be further phosphorylated to 5F-dUTP and incorporated directly into DNA during replication. Because 5-FU induced only a small number of SBS17 mutations at a dose close to its IC50, we first tested the 2′-deoxynucleoside 5F-dUridine (5F-dUrd), which should bypass RNA toxicity and channel the drug towards DNA. Repeating the mutation-accumulation experiment with 5F-dUrd from 1 nM to 1 μM, we found that the nucleoside strongly induced SBS17, being approximately 5,000-fold more potent than the base 5-FU (**Fig. 2b, c, Extended Data Fig. 4a**). We next examined whether SBS17 mutations are induced by dNTP imbalance or incorporation of 5F-dUTP into DNA.

**Fig. 2.**
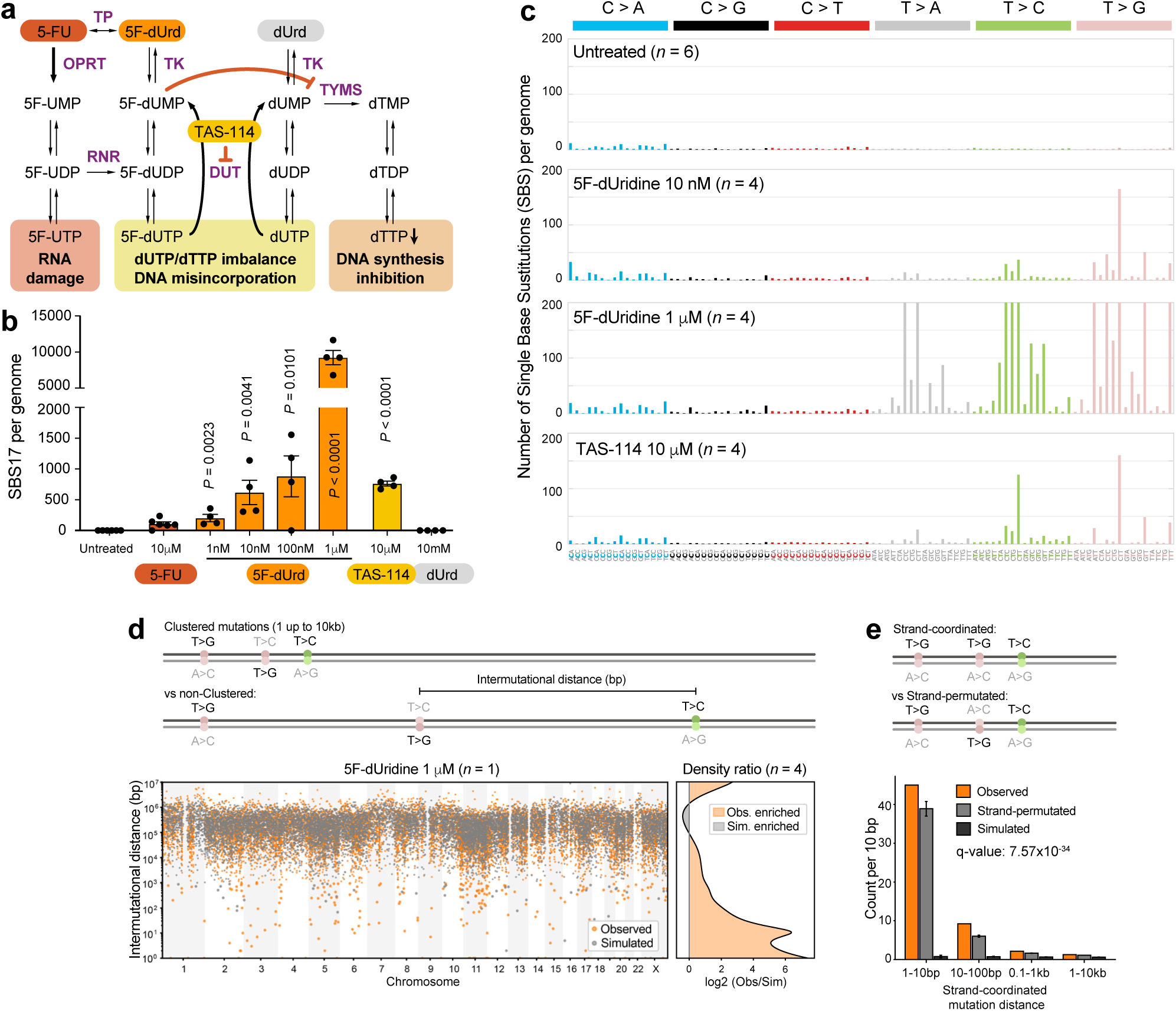
SBS17 is caused by dUTP pool imbalance. **a)** Scheme illustrating dNTP metabolism, highlighting relevant enzymes and compounds. TP: Thymidine phosphorylase; TK: Thymidine kinase; OPRT: Orotate phosphoribosyltransferase; TYMS: Thymidylate Synthase; RNR: Ribonucleotide reductase; DUT: dUTPase, deoxyuridine triphosphate pyrophosphatase. **b)** Burden of SBS17 mutations per (haploid) genome (*P* calculated by two-tailed unpaired *t* test, data shown as mean and s.e.m., *n* = 4-6). **c)** 96-classes of single-base substitutions (SBSs) considering the six mutation types but also the bases immediately 5’ and 3’ of the mutated base. Each graph represents the average mutation pattern for *n* genomes, where *n* is indicated in each panel. **d)** Left, distance between subsequent SBSs in a representative 5F-dUrd sample and a simulated dataset generated from this sample by SigProfilerSimulator. Right, log2-fold change of kernel density estimation for observed over simulated mutation distances, *n* = 4. **e)** Occurrence of strand coordinated pairs of mutations assigned to SBS17, for observed mutations, strand-permuted mutations and mutations simulated with SigProfilerSimulator. Error bar represents 95% confidence intervals, calculated as mean ± 1.96 × s.e.m. q value represents the Benjamini-Hochberg corrected p-value from a one-sided z-test against the strand-permuted null.

To imbalance the dUTP/dTTP pool by an independent mechanism, we exposed cells to TAS-114, an inhibitor of dUTPase (DUT), the enzyme that hydrolyses dUTP to dUMP to prevent its incorporation into DNA^30^ (**Fig. 2a**). Because DNA polymerases cannot distinguish dUTP from dTTP, DUT is essential to prevent uracil incorporation into DNA during replication, as established in both *E. coli* ^31^ and yeast ^32^. Remarkably, by repeating the mutation-accumulation experiment with TAS-114 (10 μM), we found that DUT inhibition also induced SBS17 mutations (**Fig. 2b,c, Extended Data Fig. 4a**). By contrast, dUridine alone was not mutagenic, consistent with efficient dUTP hydrolysis by DUT (**Fig. 2b**). Although the spectra induced by TAS-114 and 5F-dUrd both broadly resembled cancer SBS17 (cosine similarities 0.9417 and 0.9284, respectively), their trinucleotide distributions differed significantly (*P* = 1 × 10^−6^), with 5F-dUrd producing additional T>N peaks in the C<u>T</u>C context (**Extended Data Fig. 4b**). Together, these results identify dUTP/dTTP imbalance as sufficient to generate SBS17 mutations without requiring direct incorporation of 5F-dUTP into DNA.

Combined 5F-dUrd and TAS-114 treatment caused no further increase in SBS17 mutations over TAS-114 alone, probably due to extreme cytotoxicity (**Extended Data Fig. 4c**). Furthermore, we saw an increase in the number of doublet-base substitutions (DBS) and small deletions at the highest 5F-dUrd dose (1 μM), but no effect on structural variants (**Extended Data Fig. 4d**). We then explored whether the SBS17 experimental signature (∼46,000 mutations), shares topographical features with SBS17 mutations in cancer. SBS17 mutations were enriched in late-replicating and intergenic regions, both common features of many mutational signatures including SBS17 (**Extended Data Fig. 5a,b**). Additionally, SBS17 mutations showed no detectable transcriptional-strand bias, arguing against a major role for transcription-coupled damage or repair in generating the signature (**Extended Data Fig. 5b**). Replication-strand asymmetry was also not detected in our experimental system, although this feature has been reported inconsistently for SBS17 in cancer^12,33^. A representative 5F-dUrd-treated genome showed extensive spatial clustering of SBS17 mutations (**Fig. 2d**), and quantitative analysis across four independent genomes confirmed enrichment of short intermutational distances relative to simulated mutation distributions. SBS17 mutations also showed strong strand coordination (*q* = 7.57 × 10^−34^; **Fig. 2e**; **Extended Data Fig. 5c**), recapitulating features reported for SBS17 in cancer^33,34^. Importantly, such strand-coordinated clusters typically arise from a single damage event confined to one strand of DNA^35^, compatible with incorporation of uracil during DNA synthesis. Together, independent perturbations of dUTP/dTTP homeostasis converge on SBS17 mutagenesis, establishing nucleotide pool imbalance as a driver of SBS17 mutagenesis and implicating uracil misincorporation as the likely intermediate.

### The uracil glycosylase UNG drives SBS17 mutations

Uracil incorporated opposite adenine (U:A) during replication has the same coding properties as thymine and is not itself mutagenic in that context^36^. Uracil is nonetheless one of the most common non-canonical bases in DNA: the spontaneous hydrolytic deamination of cytosine to uracil generates a promutagenic U:G mispair that templates C>T mutations if it is replicated before repair^37,38^. Cells counter this with dedicated uracil-DNA glycosylases that excise uracil to initiate base excision repair (BER): the glycosylase leaves an AP site that is incised by APE1 and then filled and sealed by polymerases and ligases to complete repair^39,40^. Because this active processing converts a non-mutagenic U:A pair into an AP site intermediate, we asked whether it was the repair of uracil, rather than uracil itself, that gives rise to SBS17 mutations.

To test this, we generated HAP1 cells deficient in the primary glycosylases that act on uracil: *UNG^−^*, *SMUG1^−^* and *UNG^−^SMUG1^−^* knockouts (**Extended Data Fig. 6a, b**), and repeated the mutation-accumulation experiment with 5F-dUrd or TAS-114. The *UNG* knockout was hypersensitive to both treatments, and this was suppressed by additional loss of *SMUG1* in the double knockout line (**Extended Data Fig. 6c**), in agreement with a recent report^41^. Because UNG-deficient cells diploidised with the treatments, we performed the mutation accumulation experiment with diploid clones, 5F-dUrd (10 nM) or TAS-114 (5 μM), and 20–25X WGS. As expected, C>T mutations were increased in *UNG^−/–^*cells and further elevated in the *UNG^−/–^SMUG1^−/–^* double knockout irrespective of treatment (**Fig. 3a,b, Extended Data Fig. 6d,e**), reflecting the redundant role of these glycosylases in removing uracil arising from spontaneous cytosine deamination at C:G base pairs^42,43^. *De novo* extraction identified a C>T-rich signature assigned to SBS30 (*NTHL1* deficiency^44,45^), but its spectrum differs from SBS30; we therefore termed it SBS-UNGKO (**Fig. 3a**; **Extended Data Fig. 6f**).

**Fig. 3.**
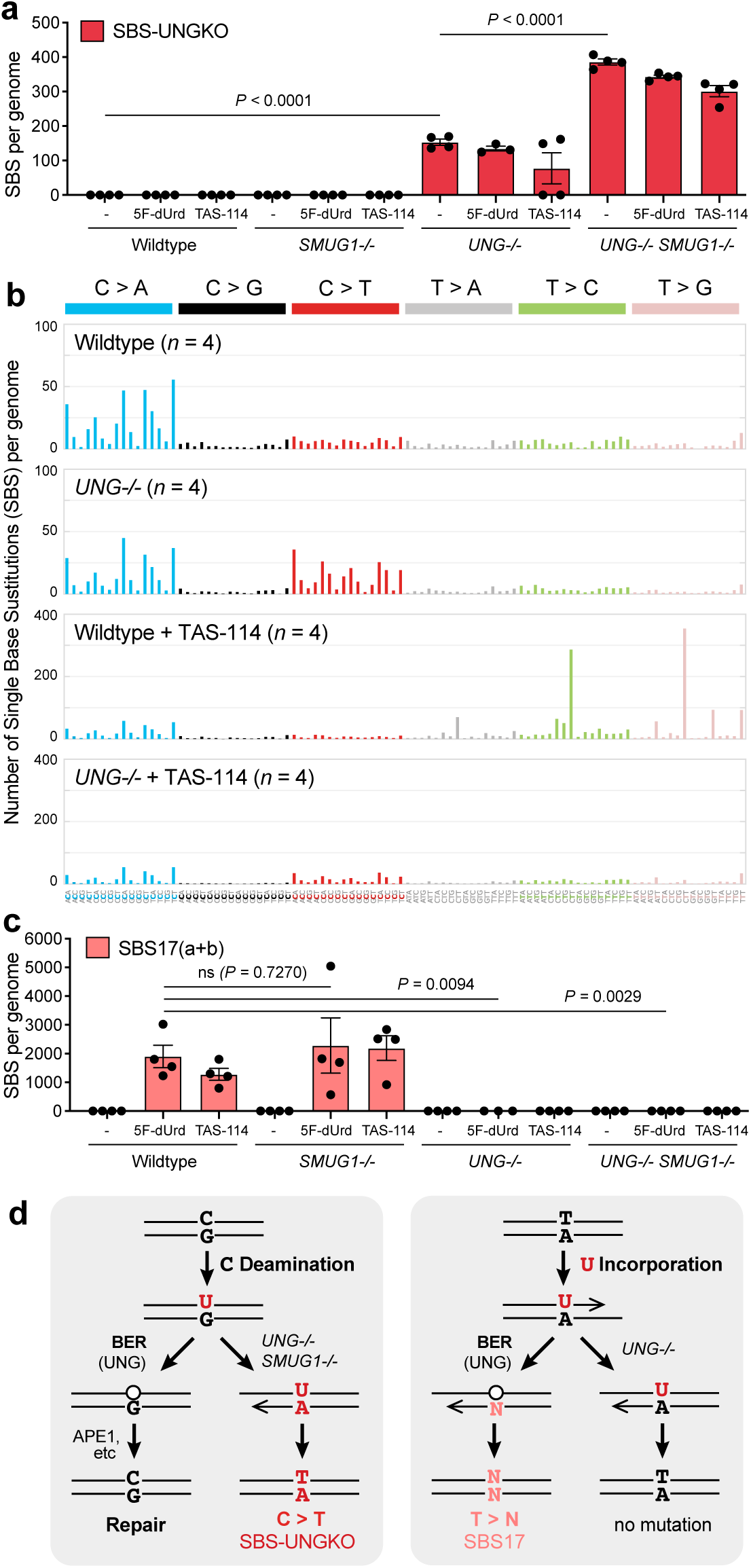
The uracil glycosylase UNG drives SBS17 mutations. **a)** Burden of SBS-UNGKO mutations (C>T) per (diploid) genome (*P* calculated by two-tailed unpaired *t* test, data shown as mean and s.e.m., *n* = 3-4). **b)** 96-classes of single-base substitutions (SBSs) considering the six mutation types but also the bases immediately 5’ and 3’ of the mutated base. Each graph represents the average mutation pattern for *n* genomes, where *n* is indicated in each panel. **c)** Burden of SBS17 mutations per (diploid) genome (*P* calculated by two-tailed unpaired *t* test, data shown as mean and s.e.m., *n* = 3-4). **d)** Cartoon illustrating the antimutagenic and mutagenic roles of UNG in processing uracil arising from cytosine deamination (U:G) versus replicative misincorporation (U:A).

Strikingly, SBS17 mutations disappeared entirely in *UNG^−/–^* and *UNG^−/–^SMUG1^−/–^* cells following either 5F-dUrd or TAS-114 treatments, whereas loss of SMUG1 alone had no effect (**Fig. 3b,c**). Uracil-DNA glycosylase activity therefore has opposing consequences depending on the lesion: excision of deamination-derived uracil (U:G) by UNG and SMUG1 protects against C>T mutations, whereas excision of uracil incorporated during DNA replication (U:A) by UNG generates SBS17 (**Fig. 3d**). This shows that SBS17 mutations do not originate from uracil itself but specifically from UNG-mediated removal of uracil from DNA.

### Translesion synthesis shapes SBS17 mutagenesis

Thus, UNG converts otherwise non-mutagenic U:A pairs into mutagenic AP site intermediates that give rise to SBS17. If not promptly repaired by BER, AP sites are non-instructional lesions with the potential to cause SBS17 mutations (as there is no hydrogen boding interface to inform complementary base-pairing). The relative heights of the T>G, T>C and T>A peaks of the SBS17 signature could reflect the probability of inserting C, G or T opposite the AP site, respectively. AP sites block replicative polymerases but can be bypassed by the translesion synthesis (TLS) machinery, a set of specialised polymerases whose larger active sites allow them to tolerate DNA lesions^46^. To test the role of TLS in the genesis of SBS17 mutations, we generated a panel of TLS-deficient mutants in HAP1 cells. This included lines disrupting each step of TLS: 1) recruitment (ubiquitylation-dead *PCNA^K^*^1c^*^4R^*, *REV1^−/–^* null, or a TLS scaffold-deletion *REV1^ΔC^*); 2) insertion (catalytic-dead *REV1^DE>GG^*or inserter polymerase knockouts *POLK^−/–^, POLH^−/–^* and *POLI^−/–^*) and 3) Polζ-mediated extension (*REV3^−/–^, REV7^−/–^)* (**Extended Data Fig. 7**). Diploid lines were treated with five cycles of TAS-114 (10 μM) and whole-genome sequenced to 20–25X coverage. Because these genetic manipulations differentially affected SBS17a and SBS17b, we quantified the two components separately (**Fig. 4a-c**).

**Fig. 4.**
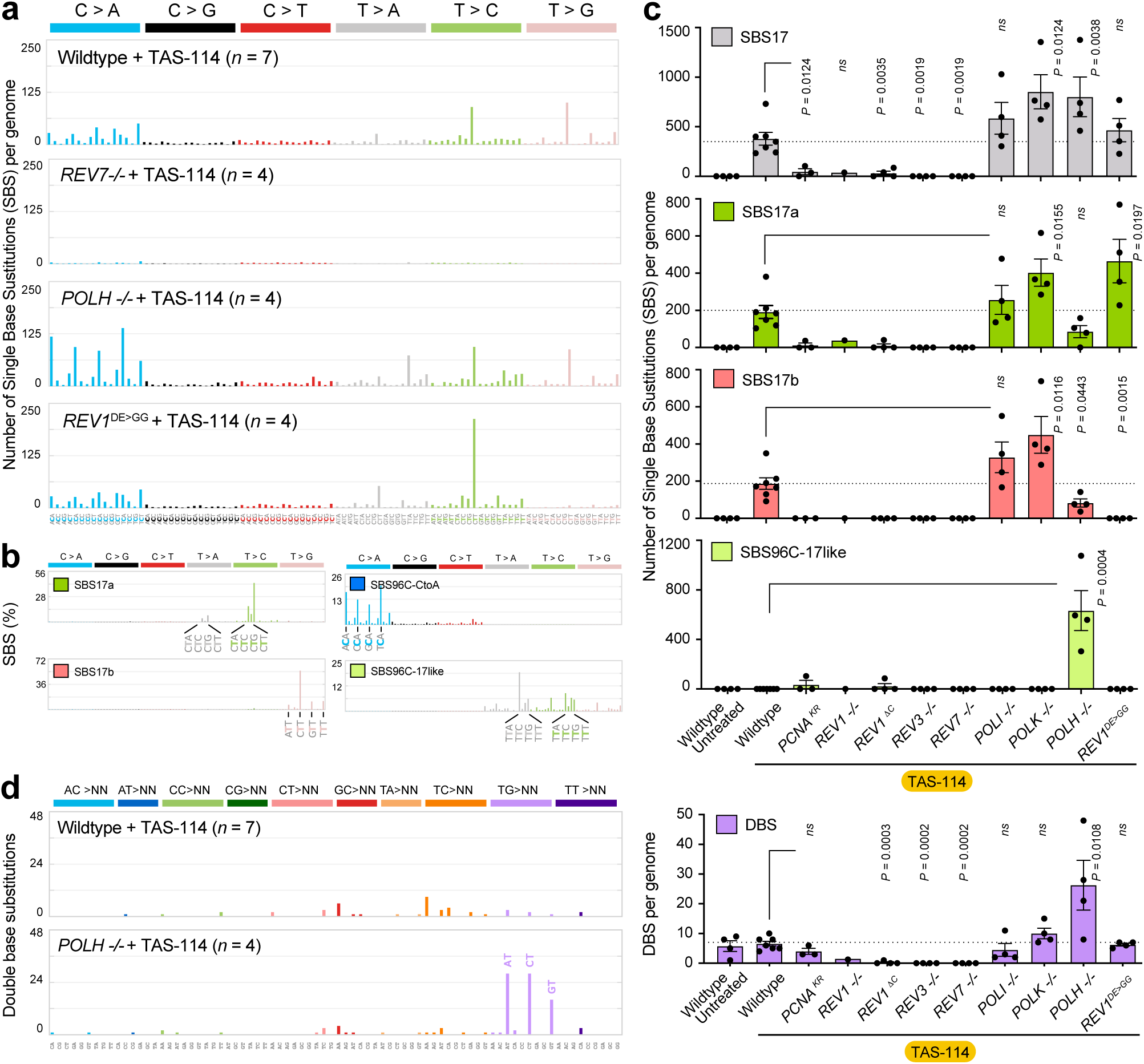
Translesion synthesis (TLS) shapes SBS17 mutagenesis. **a)** 96-classes of single-base substitutions (SBSs) considering the six mutation types but also the bases immediately 5’ and 3’ of the mutated base. Each graph represents the average mutation pattern for *n* genomes, where *n* is indicated in each panel. **b)** Pattern of known COSMIC mutational signatures and the novel signature SBS96C, split into C>A and SBS17-like components, 96-classes of SBSs considering the six mutation types but also the bases immediately 5’ and 3’ of the mutated base. **c)** Burden of total SBS17 mutations, SBS17a, SBS17b and SBS17-like mutations per (diploid) genome (*P* calculated by two-tailed unpaired *t* test, data shown as mean and s.e.m., *n* = 3-6, but *n* = 1 for *REV1-/-*). **d)** Pattern of doublet base substitutions (DBSs), following the 78-type classification from the COSMIC database. Due to the low number of DBSs per sample, the data from *n* genomes, where *n* is indicated in each panel. Right, burden of DBSs per (diploid) genome (*P* calculated by two-tailed unpaired *t* test, data shown as mean and s.e.m., *n* = 3-6, but *n* = 1 for *REV1-/-*).

We found that TAS-114 failed to induce SBS17 mutations in the *PCNA^K^*^1c^*^4R^, REV1^ΔC^*, *REV3^−/–^* and *REV7^−/–^* lines, indicating that TLS recruitment and extension by the ubPCNA-REV1-Polζ complex are essential for the generation of SBS17 mutations (**Fig. 4a,c, Extended Data Fig. 8a,b**). A single *REV1^−/–^* clone showed a similarly marked reduction, although this could not be statistically evaluated. Among the remaining mutants, loss of Polι (*POLI^−/–^*) had no effect on the burden of SBS17 mutations; and loss of Polκ (*POLK^−/–^*) caused a significant increase in total SBS17 (*P* = 0.0124), consistent with Polκ normally performing relatively error-free insertion opposite the AP site (**Fig. 4a, c**). Loss of Polη likewise increased total SBS17 (*P* = 0.0038), including SBS17b (*P* = 0.0443), but, also markedly altered the sequence context of the resulting mutations, as explored below. Most strikingly, the T>G (SBS17b) mutations were completely absent (*P* = 0.0015) in catalytic-dead REV1 cells (*REV1^DE>GG^*), indicating SBS17b is entirely dependent on the deoxycytidyl transferase activity of REV1 which inserts exclusively cytosine opposite AP sites^47,48^. Loss of SBS17b was accompanied by a concomitant increase in SBS17a mutations (T>C and T>A), consistent with insertion of G and T by back-up polymerases (*P* = 0.0197; **Fig. 4a, c**). These results therefore provide a mechanistic basis for the computational separation of SBS17 into SBS17a and SBS17b^4,6^, where each component is consequence of a different TLS insertion mechanism.

Loss of Polη (*POLH^−/–^*) yielded a novel signature (SBS96C), comprising C>A and SBS17-like T>N components. The T>N mutations resembled SBS17a (C<u>T</u>N > C<u>C/A</u>N) but with an increase in mutations in a T<u>T</u>N sequence context (*P* = 0.00066; TVD = 0.255; **Fig. 4b, Extended Data Fig. 8c**). This indicates that Polη protects against T and G misincorporations opposite AP sites at T<u>T</u>N positions. Loss of Polη additionally increased doublet-base substitutions (DBS) analogous to SBS17 (TG > NT in place of T > N, **Fig. 4d**). Out of all the new C>A (or G>T) mutations, 88% occurred in the same context as these DBS (5’N<u>C</u>A 3’ = 5’T<u>G</u>N 3’) (**Fig. 4b**). We propose DBS and C>A substitutions reflect double or single misincorporation opposite G-AP sites, and these events are normally suppressed by Polη (**Extended Data Fig. 8d**), with DBS occurring at 6% of the observed C>A events. These results place Polη as a key factor in the genesis of SBS17a mutations. Different inserter polymerases therefore provide a plausible explanation for the different sequence contexts of SBS17a (C<u>T</u>N > C<u>C/A</u>N) and SBS17b (N<u>T</u>T > N<u>G</u>T), thus the sequence context of SBS17 reflects not only the distribution of uracil-derived AP sites but also their mutagenic bypass.

Failure to bypass AP sites in TLS-deficient lines is expected to cause replication stress and potentially genomic instability. However, when we exposed the TLS mutants to a range of TAS-114 or 5F-dUrd concentrations (**Extended Data Fig. 9a**), we observed no clear hypersensitivity relative to wildtype controls, even though key mutants were hypersensitive to the bulky genotoxin 4-nitroquinoline 1-oxide (4-NQO) (**Extended Data Fig. 7**). Consistent with this, we detected no significant increase in indels or structural variants in response to TAS-114 treatment (**Extended Data Fig. 9b**). On the other hand, we detected a 1-10kb deletion signature in *REV1^ΔC^, REV3^−/–^* and *REV7^−/–^* mutants, but this was not caused by the treatment (**Extended Data Fig. 9c, d**), in agreement with a previous report^49^. These observations suggest that AP sites can be tolerated by TLS-independent pathways, potentially including template switching^50,51^. Together, these results define the division of labour among TLS components at uracil-derived AP sites and demonstrate that SBS17 substitutions arise through their mutagenic TLS bypass.

### Spontaneous SBS17 mutations are caused by the same UNG-TLS pathway

Thus far, we have used treatments to experimentally manipulate the dUTP/dTTP pool, but gastric tissue is not normally exposed to TAS-114 or 5F-dUrd. We therefore asked whether the same mechanism applies to spontaneous SBS17 mutations. We recently found that mouse liver cholangiocyte organoids spontaneously accumulate SBS17 mutations^52^, and similar observations have been reported in mouse embryonic fibroblasts (MEFs) and mouse small intestinal organoids^53,54^. To test whether this phenomenon was reproducible across cell lineages, we examined two unrelated mouse cell types, 32D (myeloid) and PPT-53631 (pancreatic), both of which spontaneously accumulated SBS17 (**Extended Data Fig. 10a**). By contrast, human cells do not accumulate SBS17 *in vitro*, including HAP1 cells in this study and previously analysed RPE-1 (ref. 49), TK6 (ref. 55), iPSCs^56^, liver and small intestinal organoids^56^, and 16 other human cell lines^57^ . Together, these data indicate that all the mouse cells sequenced to date possess a cell-intrinsic property that promotes SBS17 accumulation during *in vitro* culture.

We used mouse organoids as a model to explore whether spontaneous SBS17 mutagenesis follows the same mechanism. For this experiment, we took clonally derived liver cholangiocyte organoid lines, split them into four cultures, passaged them for four months and sequenced the bulk cultures by NanoSeq (**Fig. 5a**). To test the role of UNG excision, we drew on seminal work demonstrating the role of UNG in antibody diversification^58^, and overexpressed the bacteriophage UNG inhibitor Ugi in wildtype organoids. To test the role of TLS, we derived clonal cholangiocyte lines from *Rev1^−/–^*, *Pcna^K^*^1c^*^4R^* and *Rev7^−/–^* mice. After four months of *in vitro* culture, wildtype organoids accumulated 0.06 SBS17 mutations per Mb (approximately ∼160 mutations/genome), whereas SBS17 was strongly reduced in Ugi clone 1 (∼0.02 per Mb, *P* = 0.0520) and nearly absent in Ugi clone 2 (*P* = 0.0064), *Pcna^K^*^1c^*^4R^* (*P* = 0.0181), *Rev1^−/–^* (*P* = 0.0088) and *Rev7^−/–^* (*P* = 0.0064) organoids (**Fig. 5b, c, Extended Data Fig. 10b**). Thus, these results establish that spontaneous SBS17 mutations, like experimentally induced SBS17, arise through the mutagenic TLS bypass of UNG-derived AP sites.

**Fig. 5.**
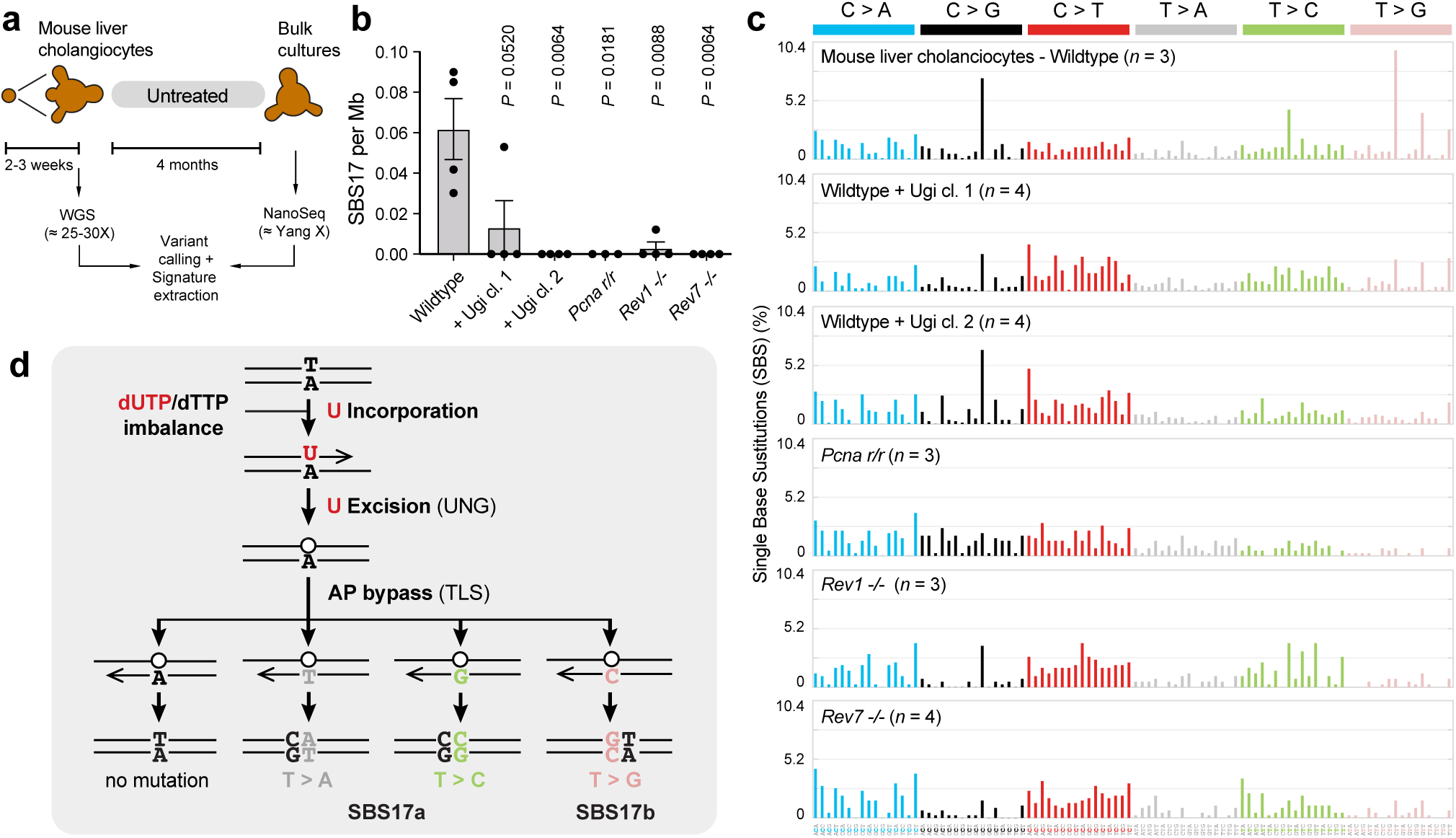
Spontaneous SBS17 mutagenesis requires UNG excision and TLS. **a)** Experimental set up for the generation of liver cholangiocyte organoid lines and assessing the mutations that arise spontaneously through *in vitro* culture by NanoSeq. **b)** Estimated burden of total SBS17 mutations per genome (*P* calculated by two-tailed unpaired *t* test, data shown as mean and s.e.m., *n* = 3-4). **c)** 96-classes of single-base substitutions (SBSs) considering the six mutation types but also the bases immediately 5’ and 3’ of the mutated base. Each graph represents the average mutation pattern for *n* bulk samples cultured in parallel, where *n* is indicated in each panel. **d)** Unified molecular mechanism of SBS17 mutagenesis.

## Discussion

Every cancer genome is a record of the mutational processes that shaped it, yet for most of the mutational signatures we can now detect, the responsible mechanism remains unknown. Understanding molecular mechanisms is a prerequisite for turning these descriptive patterns into something we can predict or prevent. Here we show that SBS17 is not a passive scar but a quantitatively important mutational process contributing mutations to cancer-driver genes and define a complete mechanistic chain for its generation.

This mechanistic chain proceeds as follows (**Fig. 5d**): dUTP/dTTP imbalance drives incorporation of uracil into DNA, UNG excises it to leave an AP site, and mutagenic bypass of the AP site by the ubPCNA-REV1-Polζ complex generates the mutation. We show that the very same pathway operates whether the trigger is a chemotherapeutic or an endogenous lesion in untreated organoids, providing a unifying mechanism for one of the most common mutational signatures in human cancer. Moreover, the catalytic activity of REV1 specifically generates SBS17b, whereas alternative TLS activities generate SBS17a, providing a biochemical basis for the computational separation of SBS17 into these two components. Strikingly, this is the very chemistry the immune system harnesses during antibody diversification^58–60^, and the same engine that underlies *kataegis,* the localised APOBEC-driven clusters of hypermutation found in many tumours^61–63^. In each case genomic uracil is converted by UNG into an AP site that is then bypassed mutagenically by TLS. In antibody diversification and *kataegis,* uracil is generated enzymatically by targeted cytosine deamination at C:G base pairs, producing U:G mismatches that can ultimately yield C>N mutations. In SBS17, the uracil is delivered through DNA synthesis, as a correctly paired U:A, and yields T>N mutations. Thus, the same fundamental sequence (uracil formation, UNG excision and mutagenic lesion bypass) can generate distinct mutation classes depending on how uracil enters DNA. SBS17 therefore reveals nucleotide-pool imbalance as a route into this mutagenic pathway that is distinct from enzymatic deamination.

By defining the molecular mechanism of SBS17, we can now frame two key questions as testable hypotheses. First, the exquisite sequence context of SBS17 could in principle be imposed at any step: uracil incorporation, UNG excision, APE1 processing, or AP site bypass. Because most AP site bypass follows the ‘A-rule’ (error-free in this context), SBS17 events represent a small mutagenic fraction of AP site bypass. Our genetic data indicate that polymerase choice contributes directly to this sequence specificity: REV1 catalytic activity drives SBS17b, whereas loss of Polη alters the sequence context of SBS17a mutations. Defining the relative contributions of lesion formation and polymerase selectivity will require biochemical reconstitution of TLS and AP site bypass across different sequence contexts. Second, the basis of the tissue specificity remains unresolved. SBS17 is absent from normal tissues^9^, but switches on during transformation in multiple tissues^7,8,11,64,65^, implying a cell-intrinsic switch. Our mouse models, in which SBS17 arises cell-intrinsically, may offer a tractable system in which to identify the underlying determinant, plausibly linked to dUTP/dTTP imbalance. Cell-extrinsic factors may also contribute to SBS17; for instance, gastric content where cytosine deamination at low luminal pH could perturb dNTP pools^66^. We predict that dUTP/dTTP ratio and genomic uracil will be elevated in SBS17-positive cancers, measuring these across human tissues would test this directly.

These findings carry therapeutic potential and illustrate how mutational signatures might be exploited more broadly. Defining the metabolic conditions that generate SBS17 may expose both preventive and therapeutic opportunities. Because SBS17 is driven by dNTP pool imbalance, it will be important to test whether restoring dUTP/dTTP homeostasis, for example through thymidine or folate supplementation, can suppress ongoing SBS17 mutagenesis. Conversely, the same dependency on dNTP imbalance could be exploited to selectively target SBS17-positive tumours. DUT inhibition has already entered clinical testing in combination with fluoropyrimidines, including TAS-114 with S-1 in advanced gastric cancer^67,68^; our finding that TAS-114 itself potently generates SBS17 raises the possibility that the mutagenic consequences of this strategy should be considered alongside its cytotoxic effects.

By resolving a long-standing signature of unknown cause into a defined enzymatic pathway, our work turns SBS17 from a descriptive mark into a mechanism that can be understood and, potentially, exploited. As the basis of other common signatures is uncovered in the same way, we envisage mutational signatures may increasingly serve as points of intervention.

## Acknowledgements

The authors would like to thank members of the Hubrecht Institute Flow Cytometry and Animal facilities for essential support. We thank Julian Sale, Martin Taylor, Gerry Crossan, KJ Patel, Puck Knipscheer and members of the Garaycoechea lab for critical reading of the manuscript. The research was supported by Dutch Cancer Society KWF Young Investigator Grant (project 12260) and ERC Starting Grant (101041308 CLOCK). The computational development reported in this manuscript have utilized the Triton Shared Computing Cluster at the San Diego Supercomputer Center of UC San Diego. This work was delivered as part of the CAUSE team supported by the Cancer Grand Challenges partnership funded by Cancer Research UK (CGCATF-2025/100021 to J.I.G.), the National Cancer Institute (1OT2CA320056 to L.B.A.) and KWF (Dutch Cancer Society).

## Author contributions

The majority of the experiments (Fig. 2-5) were carried out by S.D.C., and initial experiments (Fig.1 and Extended Data Fig. 3) carried out by N.W. and L.L.; organoid isolation and culture, S.D.C.; bioinformatic analysis, J.v.S., Y.J., L.B., bioinformatic analysis of driver mutations, A.A. L.B.A.; NanoSeq sequencing and data analysis, Y.J.; mouse husbandry and experimentation, J.W.; figure preparation, S.D.C., J.v.S., J.I.G.; study concept and design, S.D.C., J.I.G.; manuscript J.I.G. with contributions from all authors.

## Data availability

All sequencing data have been deposited at the European Nucleotide Archive (ENA) under accession code PRJEB123598. Code and data to reproduce the figures presented in this paper are available on https://github.com/GaraycoecheaGroup/SBS17_figures

## Code availability

Variant calling pipeline for whole-genome sequencing data: https://github.com/GaraycoecheaGroup/MuFASA. Statistical comparison of mutational spectra: https://github.com/GaraycoecheaGroup/MutModels.

## Methods

### Study cohorts and genomic datasets

We analysed somatic mutation data from four previously published whole-genome sequencing (WGS) resources spanning premalignant, primary, and metastatic gastrointestinal disease. No whole-exome sequencing-derived mutation catalogues were included. Primary WGS cohorts from The Cancer Genome Atlas (TCGA) comprised colorectal cancer (*n*=362), stomach cancer (*n*=418), and oesophageal adenocarcinomas (EACs; *n*=14). TCGA data were accessed through the database of Genotypes and Phenotypes (dbGaP; accession phs000178.v11.p8)^70–72^. An additional 520 primary EACs were obtained from the Mutographs project, yielding a combined primary EAC cohort of 534 tumours^73^. Metastatic WGS cohorts were obtained from the Hartwig Medical Foundation pan-cancer resource and comprised 877 colorectal cancers, 54 stomach cancers, and 177 EACs^74^.

To examine SBS17 in a premalignant setting, we additionally analysed the longitudinal Barrett’s oesophagus (BE) WGS cohort reported by Paulson et al.^75^.This dataset contains 427 longitudinal samples from 80 patients, including 40 patients with stable BE (non-progressors) and 40 who subsequently progressed to EAC (progressors)^75^.

For pan-cancer analyses of SBS17 prevalence, we additionally included WGS cohorts of primary and metastatic bladder, breast, head and neck, lung, lymphoid, and pancreatic cancers represented in TCGA and Hartwig. Driver-focused analyses were restricted to EAC, stomach cancer, colorectal cancer, and BE because other gastrointestinal malignancies had insufficient cohort sizes for robust driver analyses.

### Mutational signature assignment and genome-wide SBS17 burden

Mutational signature exposures were estimated using SigProfilerAssignment^76^ with COSMIC v3.5 SBS reference signatures^4^. GRCh38 reference signatures were used for TCGA, Mutographs, and BE samples, whereas GRCh37 signatures were used for Hartwig samples. Unless otherwise indicated, SBS17a and SBS17b activities were summed before downstream analysis and treated as a single SBS17 process. For the fluoropyrimidine-exposure analysis, SBS17a and SBS17b were analysed separately. A sample was classified as SBS17-positive when its combined SBS17a+SBS17b mutation count was greater than zero.

Genome-wide SBS17 prevalence was compared between primary and metastatic tumors using two-sided Fisher’s exact tests, with Benjamini-Hochberg false-discovery-rate (BH-FDR) correction across cancer types. For EAC, stomach, colorectal and lymphoid cancers, SBS17 burden and relative contribution were evaluated among SBS17-positive tumours. For BE, all 40 non-progressors and 40 progressors were retained; biopsy-level SBS17 counts were averaged by patient so that each patient contributed a single value. Prespecified between-group burden comparisons were assessed using two-sided Mann–Whitney U tests.

### Fluoropyrimidine exposure analysis

For the fluoropyrimidine analysis, metastatic tumours from the Hartwig cohort were classified as exposed when treatment records documented 5-fluorouracil (5-FU) or capecitabine exposure. SBS17a and SBS17b were analysed separately. Among tumours positive for the respective signature, mutation burdens were compared between exposed and unexposed groups using two-sided Mann–Whitney U tests. Relative signature contributions were summarized as medians with 95% bootstrap confidence intervals. Lymphoid cancer was reported descriptively because only one fluoropyrimidine-exposed sample was available.

### Driver gene and mutation identification

Candidate driver genes and mutations were identified using dNdScv, which estimates positive selection from synonymous and nonsynonymous mutation rates while accounting for sequence context and gene-specific mutation rates^77^. dNdScv was run in SNV-only mode separately for each primary and metastatic cancer cohort. Primary EAC samples from TCGA and Mutographs were combined before dNdScv analysis, whereas metastatic EAC was analyzed independently in Hartwig. For BE, mutations observed across biopsies from the same patient were pooled into a single patient-level mutation catalogue before dNdScv analysis, avoiding pseudoreplication of the same clonal event across biopsies.

Candidate driver SNVs were restricted to genes reaching gene-wide dNdScv significance (qallsubs_cv < 0.1). Within these genes, mutations were retained only when the estimated driver probability for their impact class was greater than zero. For an impact-class dN/dS ratio *ω* > 1, driver probability was calculated as:

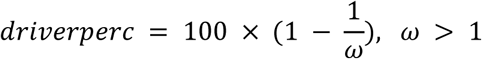

and was set to 0 when ω ≤ 1.

Candidate driver mutations were then restricted to independently validated driver genes. For colorectal cancer, stomach cancer, and EAC, genes were required to occur in the corresponding cancer type-specific IntOGen driver-gene sets (COADREAD, STAD, and ESCA, respectively)^9^. Because no directly equivalent IntOGen category exists for BE, BE driver mutations were restricted to a selected set of nine previously reported gene/transcript targets as significantly selected in this cohort^6^: *TP53*, *CDKN2A* (p16INK4a), *ARID1A*, *CDKN2A* (p14ARF), *SMARCA4*, *MUC6*, *ARID1B*, *CD5L*, and *FBXW7*. Because BE driver mutations were restricted to the previously reported BE-specific selected genes, whereas invasive cancer driver mutations were filtered using the corresponding IntOGen cancer-type driver sets, absolute driver prevalence should not be compared directly between BE and the invasive cancer cohorts.

### Attribution of individual driver mutations to mutational signatures

For each validated driver SNV, posterior signature attribution was calculated from the patient-specific signature exposure and the COSMIC SBS96 probability of the mutation’s trinucleotide context:

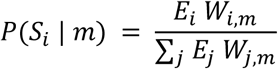

Here, *E_i_* is the exposure to signature i in the corresponding sample and *W_i,m_* is the COSMIC reference probability assigned by signature i to the SBS96 channel of mutation m; the denominator sums over signatures present in both the fitted exposure profile and the reference matrix. SBS17a and SBS17b posterior probabilities were summed to obtain a single SBS17 posterior before any downstream calculation, including determination of the highest-posterior (argmax) signature for each mutation.

For BE, mutational signature assignment was performed independently for each biopsy, whereas driver mutations were defined from patient-pooled mutation catalogues. Biopsy-level signature exposure vectors were averaged to obtain one patient-level exposure profile. This patient-level profile was then used to calculate posterior signature attribution for the corresponding pooled driver mutations. Quantifying SBS17 involvement in driver genes and mutations

SBS17 involvement in driver mutations was evaluated using two complementary measures:

<u>1) Patient-level SBS17 driver prevalence</u> was defined as the proportion of all patients in a cohort carrying at least one validated driver mutation for which merged SBS17 had the highest posterior probability among the fitted signatures. BE non-progressors were compared with progressors, and primary cancers with their metastatic counterparts, using two-sided Fisher’s exact tests. Benjamini–Hochberg correction was applied across the primary-versus-metastatic cancer comparisons.
<u>2) Gene-level SBS17 attribution</u> was defined as the mean SBS17 posterior probability among validated driver mutations affecting each gene within a cohort. Gene-specific enrichment was assessed by comparing mutations in each gene with mutations in all other validated driver genes in the same cohort using a one-sided patient-block bootstrap (2,000 resamples). BH-FDR correction was applied independently within each cohort. Selected genes with statistically enriched SBS17 attribution were additionally examined at the protein level. Driver mutations were mapped to amino-acid coordinates, recurrent genomic hotspots were defined as the same genomic position and allele observed in at least two distinct patients, and signature posterior contributions were summarized at each protein position. Major protein domains and structural regions were obtained from canonical reviewed human UniProtKB^78^.

### Cell lines

HAP1 cells (Haplogen), 32D cells (RRID:CVCL_0118) and PPT-53631 (kindly donated by Prof. Roland Rad) were grown in IMDM media (Gibco, Cat# 12440061), RPMI 1640 (Gibco, Cat#11875093) and DMEM (Gibco, Cat# 31966021) respectively. All media were supplemented with 10% fetal bovine serum (FBS; Gibco, Cat# A5256701) and 1× Penicillin-Streptomycin (PS; Gibco, Cat# 15070063). RPMI 1640 was additionally supplemented with 5 ng/mL interleukin-3 (IL-3; Gibco, Cat# 213-13). Cells were cultured at 37°C and 5% CO_2_.

### Generation of mutant cells in HAP1

Oligonucleotides containing gRNAs were hybridizes and cloned in pSpCas9(BB)-2A-GFP (px458) (Addgene, Cat# 48138) (*MTH2/NUDT18*, *MTH3/NUDT15*, *NUDT5*, *OGG1, REV7/MAD2L2*, *REV3*, *POLI, REV1^ΔC^*), or gRNA pairs in pSpCas9n(BB)-2A-GFP (PX461) (Addgene, Cat# 48140) (*MTH1/NUDT1*, *MUTYH*, *UNG*, *SMUG1*, *REV1*, *POLH*, *POLK*). To introduce point mutations in the endogenous *loci* by base editing, gRNAs for *PCNA^K^*^1c^*^4R^* and *REV1^DE>GG^* point mutants were designed as previously described^79^, and cloned into pFYF1320 EGFP Site#1(Addgene, Cat# 47511), replacing the EGFP gRNA sequence.

24 hours prior transfection 6×10^5^ cells were seeded in a 6-well plate. Transfection was performed with Xfect™ Transfection Reagent (TaKaRa, Cat# 631318) using a total of 15ng of vector. Vectors containing gRNAs for *PCNA^K^*^1c^*^4R^* and *REV1^DE>GG^* were co-transfected with the base editor plasmid pCMV-T7-ABE8e-nSpRY-P2A-EGFP (KAC1069) (Addgene, Cat#185912) kindly gifted by Maarten Geurts. 72 hours later, cells were stained with Hoechst 34580 (Molecular Probes™, Cat# 11584876) to select for DNA content.

Haploid, GFP+ cells were single cell sorted using BD FACSJazz™ Cell Sorter and clones were grown for 14 days. Afterwards, genomic DNA was isolated, and the CRISPR-targeted site was amplified by PCR using GoTaq® G2 Flexi DNA Polymerase (Promega, Cat# M7808). PCR products were screened by Sanger sequencing to characterize the genetic modifications. Clones carrying desired frameshift mutations, large deletions (in the case of *REV1^ΔC^* and *POLI*) or base substitutions (in the case of *PCNA* and *REV1*) were expanded and further validated by Western blotting, qPCR and sensitivity to DNA damaging agents as described below. A list of gRNAs and primer sequences, as well as the genetic modifications induced by CRISPR is provided in **Table 1**. A diagram of the genetic modifications in shown in **Extended Data Figs. 2, 6** and **7**.

**Table 1.**
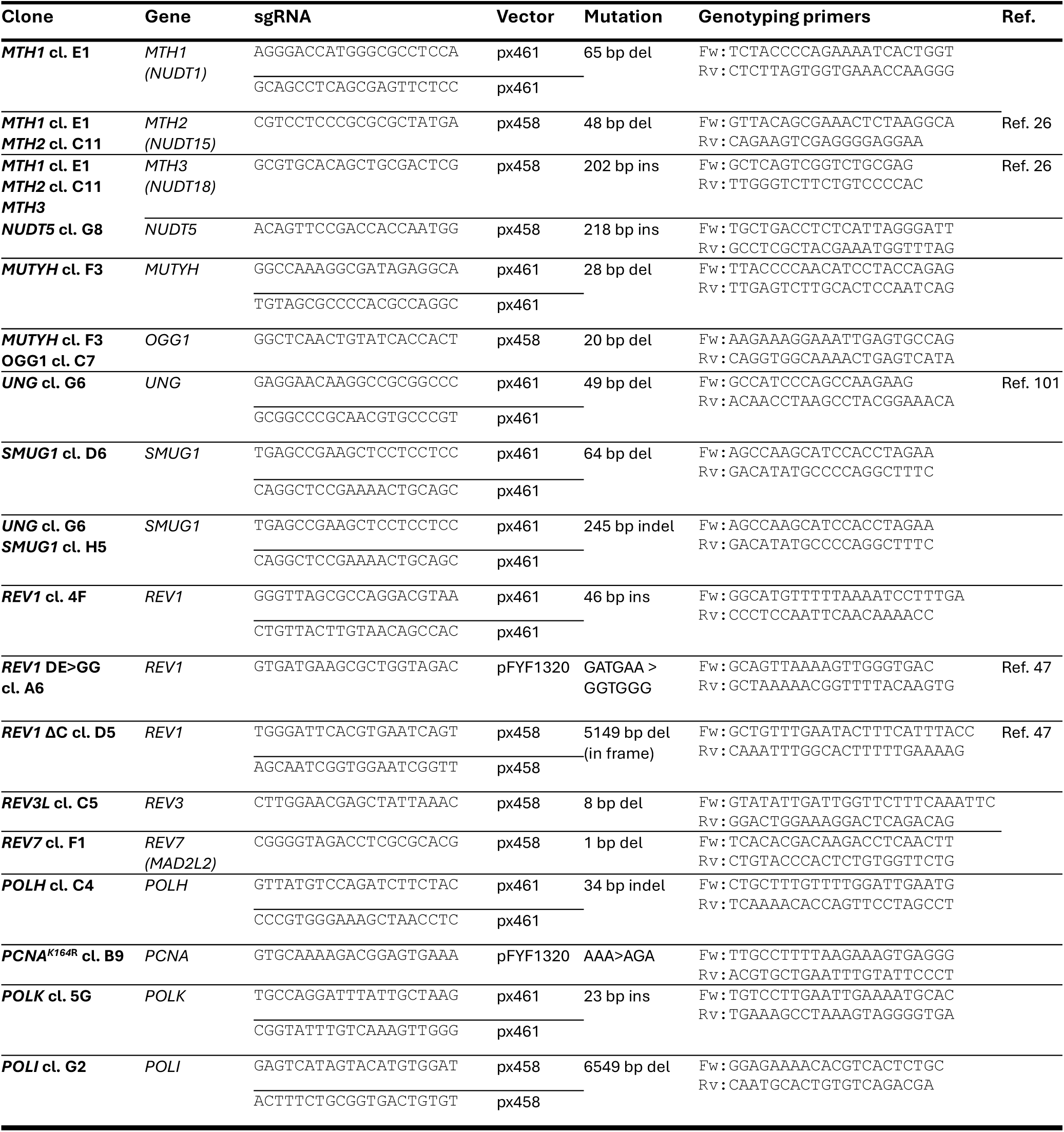
Summary of HAP1 mutant lines generated by CRISPR/Cas9.

### Western blotting

To confirm knockout of MTH1, MTH2, MTH3, and NUDT5, 1 × 10⁶ cells were harvested. To validate the *PCNA^K^*^1c^*^4R^* point mutation, we tested its inability to undergo UV-induced ubiquitination: wildtype and *PCNA^K^*^1c^*^4R^* cells were seeded in 10 cm dishes (4 × 10⁶ cells/dish) and allowed to attach overnight. Prior to irradiation, growth medium was replaced with 3 ml of PBS and cells were exposed to UV-C, with dose delivered as calibrated by a UVX radiometer. Cells received the following UV doses and corresponding exposure times: 0 J/m² (0 min), 20 J/m² (1.04 min), 40 J/m² (2.08 min), 60 J/m² (3.125 min), and 80 J/m² (4.167 min). Following irradiation, cells were then returned to growth medium and incubated for 6 h before being pelleted by centrifugation. Whole cell extracts for *MTH1*, *MTH2*, *MTH3*, *NUDT5*, *PCNA^K^*^1c^*^4R^* were prepared by resuspending cell pellets in 100µL of RIPA buffer pH 7.4 (10 mM UltraPure™ Tris Buffer (Invitrogen™, Cat#15504020), 1mM EDTA, 140 mM NaCl, 0.1% SDS, 0.5% sodium deoxycholate, 1% Triton X-100, 1 mM NaF) supplemented with protease inhibitor (Thermo Scientific™, Cat# A32953), 1U benzonase (Sigma Aldrich, Cat# E1014,), and 0.2 mM MgCl₂. Samples were incubated 5 minutes on ice, centrifuged 17000 × g for 30 minutes at 4°C and then mixed with SDS loading buffer, containing 200mM Tris-Cl, 40% glycerol, 8% SDS and 400mM DTT.

Whole cell extracts for *MUTYH*, *OGG1*, *UNG*, *SMUG1*, *POLI*, *REV1*, *REV1^DE>GG^* and *REV1^ΔC^* were instead obtained by directly mixing the cell pellets with 2× Laemmli sample buffer (40 mM Tris-HCl pH 6.8, 3.35% SDS, 16.5% glycerol, 0.005% bromophenol blue, 50 mM DTT) to a final concentration of 1 × 10⁴ cells per µL. Benzonase was added to the suspension and lysates were incubated 1 hour at room temperature.

Samples prepared in RIPA buffer were boiled at 95°C for 5 minutes, whereas samples prepared in Laemmli buffer were boiled at 70°C for 10 minutes. Proteins were resolved on NuPAGE™ Bis-Tris Mini Protein Gels, 4–12% (Thermo Scientific™, Cat# NP0321BOX) and then transferred to PVDF Membrane, 0,45µm pore size (Miliipore, Cat# IPVH00010). Membranes were blocked in 5% milk powder or 5% BSA dissolved in TBS containing 0.1% Tween-20 and incubated with primary antibodies overnight at 4°C under gentle agitation. The following primary antibodies were used: anti-β-actin (1:1000; Santa Cruz, Cat# sc-47778), anti-vinculin (1:1000; Santa Cruz, Cat# sc-73614), anti MTH1 (1:1000; Cell Signaling, Cat# D6V4O), anti-MTH2 (1:500; antibody kindly donated by Kazunari H., described in ref. 26) anti-MTH3 (1:500; Sigma A., Cat# HPA030125), anti-NUDT5 (1:500, Abcam, Cat# EPR7735), anti-MUTYH (1:500; Abnova, Cat# H00004595-M01), anti-OGG1 (1:1000; Cell Signaling, Cat# 46271), anti-UNG (Proteintech, Cat# 12394-1-AP), anti-SMUG1 (Santa Cruz, Cat# sc-514343), anti-REV1 (1:1000; Santa Cuz, Cat# sc-393022), anti-PCNA (1:5000; Santa Cruz, Cat# sc-56 HRP), anti-PCNA-ub (1:1000; Cell Signaling, Cat# D5C7P), anti-POLI (1:1000; Abcam, Cat# ab157244).

Afterwards, membranes were washed 3 times in TBS containing 0.1% Tween-20 and incubated for 1 hour with HRP-conjugated secondary antibodies at room temperature. The following secondary antibodies were used: goat anti-rabbit IgG (1:10,000; Jackson, Cat# 111-035-003), and rabbit anti-mouse IgG (1:10,000; Jackson, Cat# 315-035-003). Finally, detection was performed using SuperSignal West Maximum Sensitivity substrate (Thermo Fisher Scientific, Cat# 34096), and signals were acquired with Amersham IQ800 system (Cytiva).

### Growth inhibition assay

Growth inhibition was determined using the CellTiter-Glo 2.0 Luminescent Cell Viability Assay (Promega, Cat# G9242). Cells were seeded at a density of 1500 cells per well in clear-bottom white 96-well plates (Costar, Cat#3610) and exposed to increasing concentrations of the indicated compounds, prepared by serial 1:2 or 1:10 dilutions. Cells were incubated for 72 hours at 37°C before the culture medium was removed. 50 µL of CellTiter-Glo 2.0 reagent, diluted 1:1 with culture medium, was then added to each well. Plates were shaken on an orbital mixer to ensure complete cell lysis and incubated for 10 minutes at room temperature to allow the luminescent signal to stabilise. Luminescence was subsequently measured using a plate reader, and cell viability was calculated following blank subtraction and expressed relative to untreated control cells.

### Quantitative PCR

For validation of *POLH* and *POLK* lines, total RNA was extracted from 5×10^6^ cells using NucleoSpin RNA kit (Macherey-Nagel). First-strand cDNA was synthesized from 500 ng of total RNA using the SuperScript III First-Strand Synthesis kit (Invitrogen, Cat# 18080051) according to the manufacturer’s instructions. Briefly, RNA was combined with 1 μl anchored oligo(dT)22 (50 μM) (or 50–250 ng random primers) and 1 μl 10 mM dNTP mix in a total volume of 14 μl, heated to 65°C for 5 min, and immediately chilled on ice. Reverse transcription was performed by adding 4 μl 5× First-Strand Buffer, 1 μl 0.1 M DTT, 1 μl RNaseOUT Recombinant RNase Inhibitor (40 U/μl; Invitrogen, Cat# 10777019), and 1 μl SuperScript III Reverse Transcriptase (200 U/μl) to each reaction (final volume 20 μl). Samples were incubated at 50°C for 30– 60 min, and the reaction was inactivated by heating at 70°C for 15 min. Synthesized cDNA was diluted to >25 ng/μl prior to use in quantitative PCR (qPCR).

qPCR was performed using FastStart Universal SYBR Green Master (ROX) (Roche, Cat# 4913850001) on a CFX Duet Real-Time PCR System, using gene-specific primers (POLK_Fw: AGCCATGCCAGGATTTATTG; POLK_Rv: GGATCGTTCATGCTCACTCA; POLH_Fw: AGTTCGTGAGTCCCGTGGG; POLH_Rv: GCTTGGCAACAAGTCTGCC; GAPDH_FW: GAAGGTGAAGGTCGGAGTC; GAPDH_RV: GAAGATGGTGATGGGATTTC). Reactions were performed in quadruplicate. Cycling conditions were as follows: initial denaturation/enzyme activation at 95°C for 10 min, followed by 40 cycles of 95°C for 15 s and 60°C for 60 s, with a melt-curve analysis performed at the end of the run. Relative gene expression was calculated using the 2^-ΔΔCt method, normalized to the reference gene (*GAPDH*).

### Mutation accumulation experiments

A single parental clone was established by single-cell sorting of HAP1 cells using either the BD FACSJazz™ Cell Sorter or the CytoFLEX SRT Cell Sorter, followed by a first clonal expansion period of approximately 14 days. In particular, for the mutation accumulation experiments shown in **Fig. 1** and **Fig. 2**, cells were stained with Hoechst 34580, and haploid cells were sorted. For the experiments shown in **Fig. 3** and **Fig. 4**, diploid cells were isolated instead.

The parental clone was subsequently divided into six aliquots. One aliquot was cryopreserved, one was collected for genomic DNA extraction to serve as the ancestral reference sample, and the remaining four aliquots were used to establish four independent mutation accumulation subcultures.

For HAP1 experiments, 1×10^4^ and 2×10^5^ cells were seeded per subculture for untreated and treated conditions respectively, and 24 hours later the indicated compounds were added. Cells were treated for 72 hours, followed by a 48-hour recovery period before reseeding. This treatment–recovery cycle was repeated five consecutive times, with each subculture propagated independently throughout the experiment. Similarly, 32D and PPT-53631 clonal lines were split into four subcultures and passaged during three and four weeks respectively, to allow spontaneous mutations to accumulate.

Finally, cells from each subculture were subjected to a second round of single-cell sorting and clonal expansion to obtain enough material for genomic DNA extraction (approximately 2-3 weeks), which was performed using QIAamp DNA Mini Kit (QIAGEN, Cat# 51306). Throughout the entirety of the mutation accumulation experiments, cells were maintained at 37 °C in a low oxygen incubator (5% O₂) to minimize the accumulation of culture-associated oxidative DNA damage and the corresponding *in vitro* mutational signature SBS18, with the exception of the experiment shown in **Extended data Fig. 3e**, which was performed also under 21% O₂.

### Chemicals

Growth inhibition assays and mutation accumulation experiments were performed using the following compounds. 5-Fluorouracil (5-FU; Sigma-Aldrich, Cat# F6627) and 5-Fluoro-2′-deoxyuridine (5F-dUrd; Sigma-Aldrich, Cat# F0503) were dissolved in DMSO to stock concentrations of 0.5 M and 0.25 M, respectively, and stored at 4 °C for up to 4 months (5-FU) or up to 14 days (5F-dUrd). TAS-114 (MedChemExpress, Cat# HY-124062) was dissolved in DMSO to a 10 mM stock and stored at −80 °C for up to 6 months. 2′-Deoxyuridine (Sigma-Aldrich, Cat# D5412) was dissolved directly in growth medium to a 0.5 M stock and stored at −20 °C for up to 1 month. 4-Nitroquinoline N-oxide (4NQO; Sigma-Aldrich, Cat# N8141) was dissolved in DMSO to a stock concentration of 100mM and stored at −20 °C. All stocks were diluted to the indicated working concentrations in growth medium immediately prior to use.

### Whole genome sequencing and variant calling

Library preparation and sequencing was performed by Novogen. The 2×150 bp paired-end sequencing was performed on the Illumina NovaSeq 6000 and NovaSeq X Plus platforms with a minimum coverage of 20X for diploid and 10X for haploid genomes. Read processing, mapping, consensus variant calling and filtering was performed using our MuFaSa pipeline v1.1 (https://github.com/GaraycoecheaGroup/MuFASA). Read processing, mapping and SNV and indel variant calling was performed as in Jiang *et al.*^52^. Briefly, reads were mapped with BWA-MEM^80^, using default settings, after which consensus variant calling was performed, taking the intersect of Strelka v2.9.10 (ref. 81) and GATK Mutect2 v4.5.0 (ref. 82). SNV filtering was performed using FINGS v1.7.2 (ref. 83) with default settings and a max variant allele frequency (VAF) in the normal of 0.01. Indels were filtered with a required mapping quality > 50, read depth >10 for diploid and >5 for haploid genomes respectively. VAF filters were applied to both snvs and indels, excluding variants with a VAF < 0.3 or > 0.7 for diploid, and variants with a VAF < 0.9 for haploid genomes. Lastly, variants present in samples from the same parental clone were discarded to eliminate potential germline variants. Additionally, structural variants were identified using consensus variant calling with manta v1.6.0 (ref. 84) and gridds v2.13.2 (ref. 85) and filtered with gridds_somatic_filter using default settings.

### Mutational signature analysis

Mutational signature analysis was performed per experiment using the SigProfiler Toolkit: MatrixGenerator v1.3.6 (ref. 86) Was used to generate count tables for single nucleotide variants, doublet-base substitutions, indels and structural variants. The resulting SBS 96-class count tables were further analysed by SigProfilerExtractor v1.2.6 (ref. 87), after which SigProfilerAssignment v1.1.1 (ref. 76) was used to assign *de novo* extracted signatures to cosmic v3.5 SBS signatures, excluding artifact and tobacco related signatures. The resulting per-sample signature assignments were used as quantification of the contribution of each signature to the mutational burden of a sample. In case of poor reconstruction of the *de novo* signatures by existing cosmic signatures, either a *de novo* extracted, or an experimentally determined signature was included in the signature assignment used for quantification.

### Topography analysis of mutational signatures

Topography analaysis was performed using the SigProfilerTopography software v1.0 (ref. 88), using default parameters and n=25 simulations to generate a null distribution. Transcription strand bias was determined using default parameters. Replication strand bias and replication timing analysis were performed using a high-resolution HAP1 Repli-Seq dataset from Klein *et al*^89^. Analysis of strand-coordinated mutation clustering was performed using a customized version of the processivity analysis from SigProfilerTopography. Firstly, since the SBS17 spectrum encompasses multiple substitutions (T>N), the analysis was adjusted to consider any set of phased and clustered T>N mutations assigned to either SBS17a or SBS17b a strand-coordinated cluster. Secondly, as we observed increased spatial clustering of mutations compared to the simulated null, we asked whether the apparent enrichment of strand-coordinated mutations compared to the simulated data could be the consequence of this general clustering rather than true strand-coordinated mutagenesis. If an observed enrichment is purely a side-effect of general spatial clustering, permuting the strand information of observed mutations should not affect the degree of enrichment. Thus, we generated an alternative strand-permuted null for strand-coordinated clustering. Enrichment and significance were determined as it is using the simulated null in the SigProfilerTopography software.

### Statistical comparison of mutational spectra

To assess whether two experimental conditions induce distinct mutational spectra, we developed a parametric bootstrap test. Observed mutations were classified by their trinucleotide sequence context, resulting in per-sample count vectors. To estimate the mutations induced by a treatment or gene deficiency, the mean background mutation counts observed in the control condition are subtracted from the count vector of each sample, setting the values to min zero and rounding to whole numbers. Under the null hypothesis the two groups share the same trinucleotide frequencies, modelled as a Dirichlet-multinomial; a multinomial with a concentration parameter to account for variability between biological replicates beyond multinomial sampling noise^90^. The null distribution fits a proportion vector common to both groups and a separate concentration parameter per sample group, to test for a difference in mean mutation frequencies, not a difference in variability. As a test statistic we use the moderated studentized maximum between difference between the mean proportion vector for each group, such that a significant value indicates at least one mutation category differs more than expected under the null, while controlling for testing multiple categories and being sceptical of categories with very small standard errors^91,92^. To obtain the null distribution, a parametric bootstrap is performed, simulating 1M datasets from the fitted null, matching the observed sample numbers and their mutational burdens. An adapted version of this method is used to compare observed mutational frequencies to a provided reference set of proportions such as those from a known cosmic signature. In this case, a single Dirichlet-Multinomial is fit with a fixed mean set to the provided proportions and the concentration parameter estimated from the observed data, using the maximum difference between the mean observed proportions and the reference as the test statistic. These methods have been made available in the MutModels software (https://github.com/joerivstrien/MutModels).

### Mice

All animal experiments were performed after institutional review by the Animal Ethics Committee of the Royal Netherlands Academy of Arts and Sciences (KNAW) with project license of AVD8010020198847. The *Rev1^tm1Ndew^* (MGI 3701945, C57BL/6J) mice were described previously and a kind gift from Niels de Wind^93^. The *Pcna^tm1Jcbs^* (MGI 3761720, C57BL/6J) mice were described previously and a kind gift from Heinz Jakobs^94^. The *Rev7/Mad2l2^tm^*^1a^(EUCOMM)*^Wtsi^* (MGI 4432091) were acquired from EUCOMM, reported embryonic previously lethal in a C57BL/6J^95^, backcrossed to the 129S4/SvJaeJ for 5 generations, then used to generate *Rev7/Mad2l2^tm^*^1a^(EUCOMM)*^Wtsi^* mice in a C57BL/6N x 129S4/SvJaeJ F1 background. We used 10-month-old mice for organoid isolation. All mice were maintained and housed under standard conditions, with ambient temperature 20–23 °C and humidity between 50–60%, *ad libitum* food and water, and on a 12-h light–dark cycle.

### Organoid culture

Primary liver cholangiocytes were isolated using a protocol previously described in by Broutier *et al*^96^. In brief, liver tissue was minced and digested in wash buffer containing 125 μg/mL collagenase (Sigma-Aldrich, Cat# C9407), 125 μg/mL dispase II (Thermo Fisher Scientific, Cat# 17105041), and 0.1 mg/mL DNase I (Sigma-Aldrich, Cat# DN25). The wash buffer consisted of DMEM (Thermo Fisher Scientific, Cat# 31966021) supplemented with 2% fetal bovine serum and PS. The resulting biliary duct fragments and surrounding stromal tissue were further dissociated into a single-cell suspension using 7× TrypLE (Gibco, Cat# A1217701). Cells were then incubated in 2% FCS containing fluorophore-conjugated antibodies against EpCAM/CD326 (6:100; eBioscience, Cat# 17-5791-82), CD45 (1:50; BioLegend, Cat# 304006), CD31 (1:50; BioLegend, Cat# 102407), and TER-119 (1:50; BioLegend Cat# 116201). Cholangiocytes were identified as EpCAM-positive and negative for CD31, CD45, and TER-119, sorted using BD Influx™ Cell Sorter and plated in serial dilutions in BME (RCD Systems, Cat# 3533-010-02). Purified cholangiocytes were then clonally expanded. Cells were maintained in Advanced DMEM/F12 (Gibco, Cat# 12634010) supplemented with 10 mM HEPES, 1× GlutaMAX, and 100 U/mL PS as the basal medium. Growth medium was further supplemented with B27 (Gibco, Cat# 12587010), 1 μM N-acetylcysteine (Sigma-Aldrich, Cat# A0737), 10 nM gastrin (Sigma-Aldrich, Cat# G9145), 50 ng/mL mouse EGF (PeproTech, Cat# AF-100-15), 50 ng/mL recombinant human HGF (PeproTech, Cat# 100-39-100UG), 100 ng/mL FGF-10 (Bio-Techne RCD, Cat# 345-FG) 1% R-spondin 3-conditioned medium, 10 mM nicotinamide (Sigma-Aldrich, Cat# N0636), and 10 μM ROCK inhibitor (ROCKi; AbMole, Cat# M1817). To promote initial organoid establishment, cultures additionally received 150 ng/mL Noggin (IPA) and 27 ng/mL Wnt surrogate (IPA) during the first four days after seeding.

For the mutation accumulation experiment, following clonal expansion, the parental cholangiocyte clone was split into five aliquots. One aliquot was cryopreserved, while the remaining four were kept in culture as independent mutation accumulation subcultures. To allow the mutations to accumulate spontaneously, subcultures were propagated for four months, at 37 °C in a low oxygen incubator (5% O₂). During this period, organoids were passaged weekly by both mechanical and enzymatic digestion with TrypLE (Gibco, Cat# 12605010), followed by TrypLE inactivation with wash buffer and centrifugation at 300g for 5 minutes at 4°C. Cell pellets were then resuspended in BME and replated in fresh culture medium. At the end of 4 months, genomic DNA was extracted and bulk cultures were sequenced by NanoSeq.

### Ugi overexpression in cholangiocyte organoids

A PiggyBac-based transposon plasmid containing an IRES sequence and hygromycin resistance cassette (kindly gifted by Delilah Hendriks^97^) was digested with XhoI/NotI restriction enzymes. The EGFP-T2A sequence, amplified using primers EGFP_Fw (TCATTTTGGCAAAGAATTCCACCatggtgagcaagggcgagg) and EGFP_T2A_Rv (tgggccaggattctcctcgacgtcaccgcatgttagcagacttcctctgccctccttgtacagctcgtccatg) was fused by PCR to a UGI-NLS sequence amplified from pBT280 (Addgene, #122610) using primers T2A_UGI_Fw (tcgaggagaatcctggcccaAgcaccaacctgtctgacatc) and UGI_NLS_Rv (CGATATCAAGCTTATCGAGCttagactttcctcttcttcttg), with Phusion High-Fidelity DNA Polymerase (NEB, #M0530). The resulting EGFP-T2A-UGI-NLS fragment was inserted into the digested vector using the Gibson Assembly Cloning Kit (NEB, Cat# E5510S). The final construct was verified by whole-plasmid sequencing (Plasmidsaurus).

Wildtype liver cholangiocytes organoids were transfected as previously described^98^. One day prior to electroporation, organoids were cultured in PS-free medium supplemented with Wnt surrogate, 1.25% DMSO (MERK, Cat# 41639), and 3 μM CHIR99021. On the day of electroporation, organoids were dissociated into clumps of 3–5 cells using dispase, TrypLE containing 10 μM ROCKi and DNase, combined with mechanical disruption. Cells were washed in ice-cold DMEM and OptiMEM (Gibco, Cat# 31985070) (both containing ROCKi) and resuspended in BTXpress buffer (BTX, Cat #45-0802). 7.2 μg of the PiggyBac transposon plasmid comprising EGFP-T2A-UGI-NLS and 2.8 μg of PiggyBac transposase plasmid (kindly gifted by Delilah Hendriks) were added to the mixture and transferred to an electroporation cuvette. Electroporation was carried out using a NEPA21 electroporator (poring pulse: 175 V, 5.0 ms, 2 pulses; transfer pulse: 20 V, 50 ms, 5 pulses). Following electroporation, organoid medium containing ROCKi was added, and cells were allowed to recover at room temperature for 30 min before being pelleted, resuspended in BME, and plated. 7 days post-electroporation, organoids were dissociated into single cells using TrypLE and mechanical disruption, plated in BME as a serial dilution, and selected with 100 μg/mL hygromycin. Selection was continued until all organoids on the untransfected negative control plate had died, after which surviving clones were picked. Stable integration of the plasmid was confirmed by detection of EGFP expression using BD LSRFortessa™ Cell Analyzer.

### Restriction-enzyme NanoSeq library preparation and data preprocessing

Restriction-enzyme NanoSeq libraries were prepared following the protocol described previously^99^, with minor modifications. Briefly, on-bead DNA fragmentation was performed using HpyCH4V (NEB, Cat# R0620) with 300 ng of genomic DNA from each sample as input. For matched normal samples, 100 ng of genomic DNA isolated from the untreated bulk culture was subjected to whole-genome sequencing (WGS) at a minimum coverage of 30x. NanoSeq libraries and matched normal WGS libraries were sequenced using 2 x 150 bp paired-end reads on a NovaSeq X platform by Novogene.

Data preprocessing was performed as described by Abascal *et al* ^99^. Briefly, sequencing reads were aligned to the mouse reference genome GRCm39 using BWA-MEM. The resulting alignments were sorted using biobambam2, as previously described^100^.

### NanoSeq variant calling

Variant calling was performed as previously described^99^. Briefly, NanoSeq requires a matched normal sample to remove germline variants. Matched normal samples were generated by WGS from the untreated bulk organoids. Candidate variants were required to fulfil the following criteria: (1) each read bundle contained at least two reads derived from each of the two original DNA strands; (2) consensus base quality scores were ≥60; (3) the difference between the primary (AS) and secondary (XS) alignment scores was >50 to retain only unambiguously mapped read pairs; (4) the average number of mismatches per read bundle was ≤2 in both the sample and matched normal; (5) the maximum number of 5’ clips needed to be 0; (6) the maximum number of improper read pairs needed to be 0; (7) base calls within the 8 bp at both the 5′ and 3′ ends of reads were excluded from analysis; (8) for SNV calling, read bundles containing indels were excluded; (9) the matched normal contained minimum 15 reads per strand at the given sites; (10) the variant allele frequency in the matched normal did not exceed 0.01; and (11) the site did not overlap a common SNP. Finally, SNVs detected in moe than one library were excluded prior to downstream analysis.

### Statistical analysis

All data were presented as mean +/- s.e.m or 95% confidence interval, as indicated in the figure legend, with individual data points shown where feasible. Mutational burdens were compared by two-tailed unpaired *t*-tests. Strand asymmetries were determined with Fisher’s exact tests corrected for multiple testing using the Benjamini-Hochberg procedure. Strand-coordinated mutagenesis was tested using a one-tailed z-test against a strand-permuted null. Statistical comparisons of mutational spectra were performed with FWER-corrected two-sided parametric bootstrap tests, described in detail in the relevant methods section. For the analysis of mutations in gastrointestinal cancers Fisher’s exact tests and Mann-whitney U tests were performed, followed by multiple testing corrections using the Benjamini-Hochberg procedure. Enrichment of driver mutations was assessed by one-sided patient-block bootstrap.

---

**Extended Data Fig. 1.**
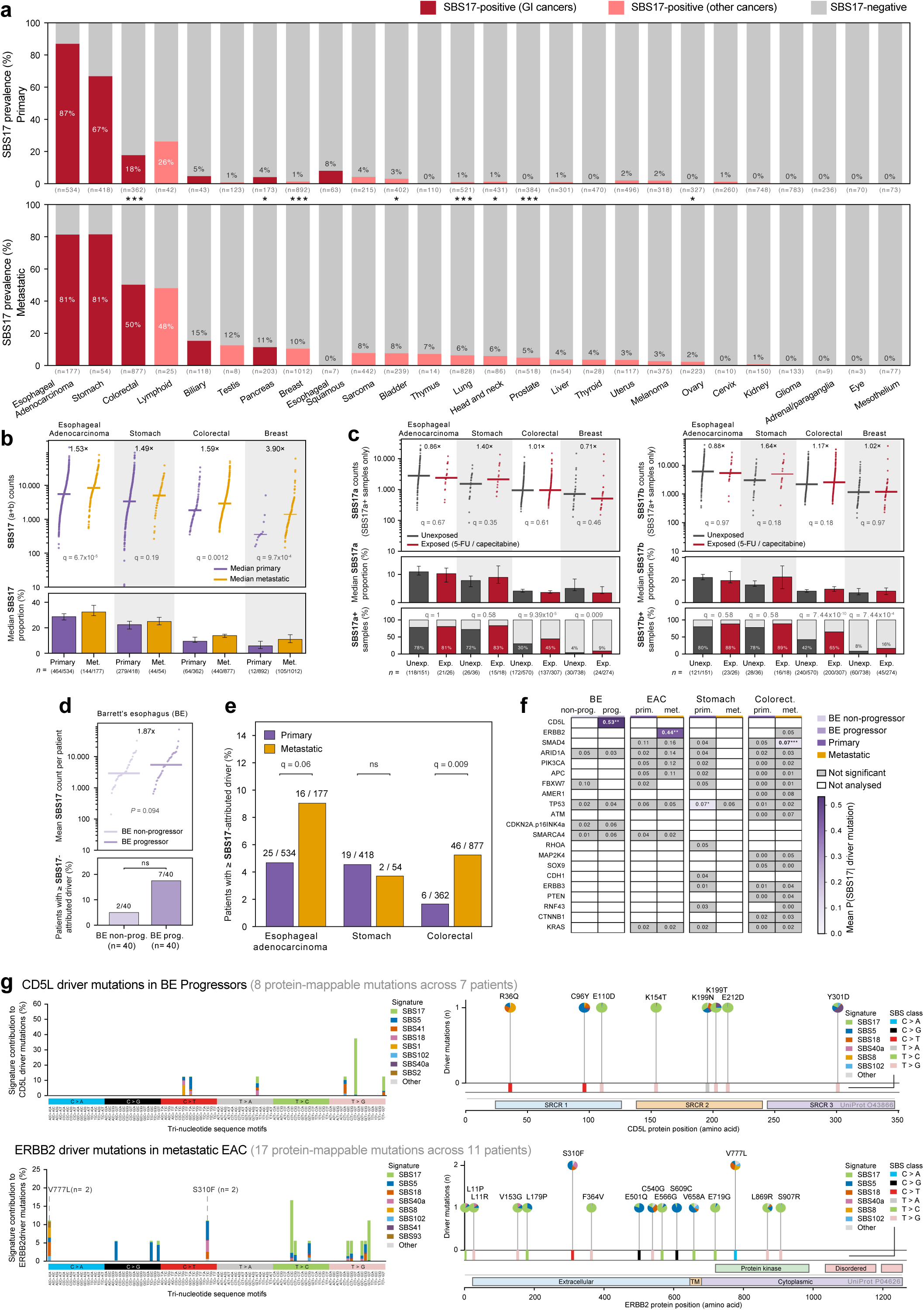
SBS17 prevalence, burden and driver attribution in gastrointestinal cancers. **a)** SBS17 prevalence across primary tumours (top) from TCGA and Mutographs and metastatic tumours (bottom) from the Hartwig Medical Foundation cohort. Bars show the percentage of SBS17-positive samples (red, gastrointestinal cancers; pink, other cancers) and SBS17-negative samples (*q < 0.05, **q < 0.01, ***q < 0.001; two-sided Fisher’s exact tests with Benjamini–Hochberg correction). **b)** SBS17 burden and relative contribution among SBS17-positive oesophageal adenocarcinoma, stomach, colorectal and breast tumours. Top, SBS17 mutation counts; horizontal lines indicate medians, and fold changes denote the metastatic-to-primary median ratios. Bottom, median percentage of substitutions attributed to SBS17; error bars indicate 95% bootstrap confidence intervals. SBS17 denotes SBS17a + SBS17b. q values were calculated using two-sided Mann–Whitney U tests with Benjamini–Hochberg correction **c)** SBS17a (left) and SBS17b (right) in metastatic tumours from patients with or without documented exposure to 5-fluorouracil (5-FU) or capecitabine. Top, mutation counts among signature-positive samples; horizontal lines indicate medians, and fold changes denote the exposed-to-unexposed median ratios. Middle, median percentages of substitutions attributed to the indicated signature; error bars indicate 95% bootstrap confidence intervals. Bottom, percentages of samples with detectable SBS17a or SBS17b. For the top panels, q values were calculated using two-sided Mann–Whitney U tests; for the bottom panels, q values were calculated using two-sided Fisher’s exact tests. All *P* values were adjusted using the Benjamini–Hochberg method. **d)** SBS17 in Barrett’s oesophagus non-progressors and progressors (n = 40 each). Left, patient-level burden averaged across longitudinal biopsies (*P*, two-sided Mann–Whitney U test). Right, patients carrying ≥1 SBS17-attributed driver (*P*, two-sided Fisher’s exact test). **e)** Patients carrying ≥1 SBS17-attributed driver in primary and metastatic oesophageal, stomach and colorectal cancers; q values are Benjamini–Hochberg-corrected Fisher’s exact-test P values. **f)** Mean posterior *P*(SBS17 | driver mutation) by gene and cohort; grey cells indicate no significance and blank cells indicate gene–cohort combinations that were not analyzed. Enrichment versus other driver mutations in the same cohort was assessed by one-sided patient-block bootstrap; *q < 0.05, **q < 0.01. **g)** Protein-level distribution of *CD5L* and *ERBB2* driver mutations and posterior signature attribution of their corresponding trinucleotide contexts.

**Extended Data Fig. 2.**
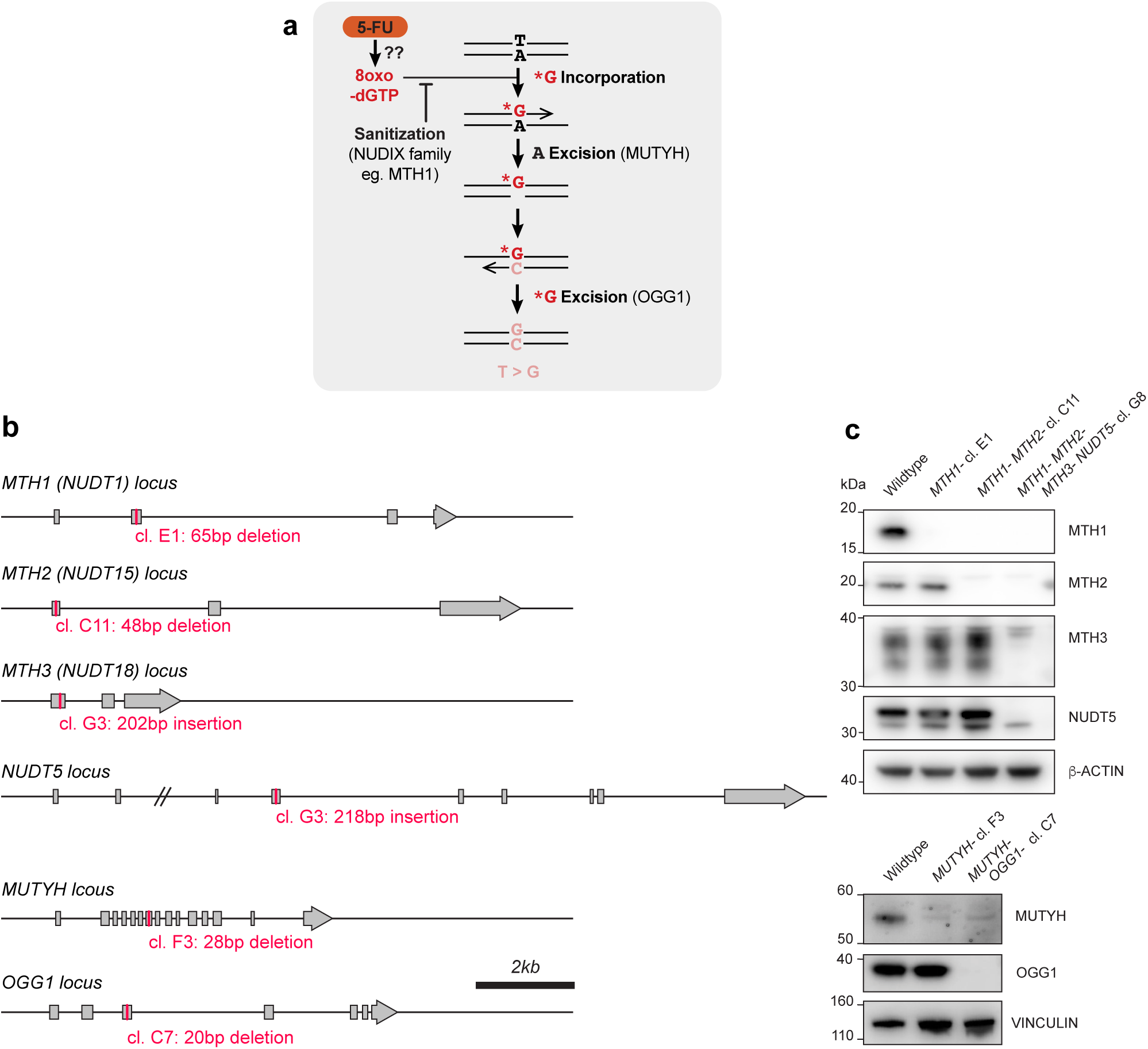
Generation and validation of mutants linked to 8-oxo-guanine metabolism. **a)** Scheme for the potential generation of T>G mutations driven by oxidation of the dGTP pool. **b)** Schematic cartoon of the human *loci*, targeted exons and introduced genetic modifications. See **Table 1** for sequence information and additional details. **c)** Validation of putative knockout lines by Western blot.

**Extended Data Fig. 3.**
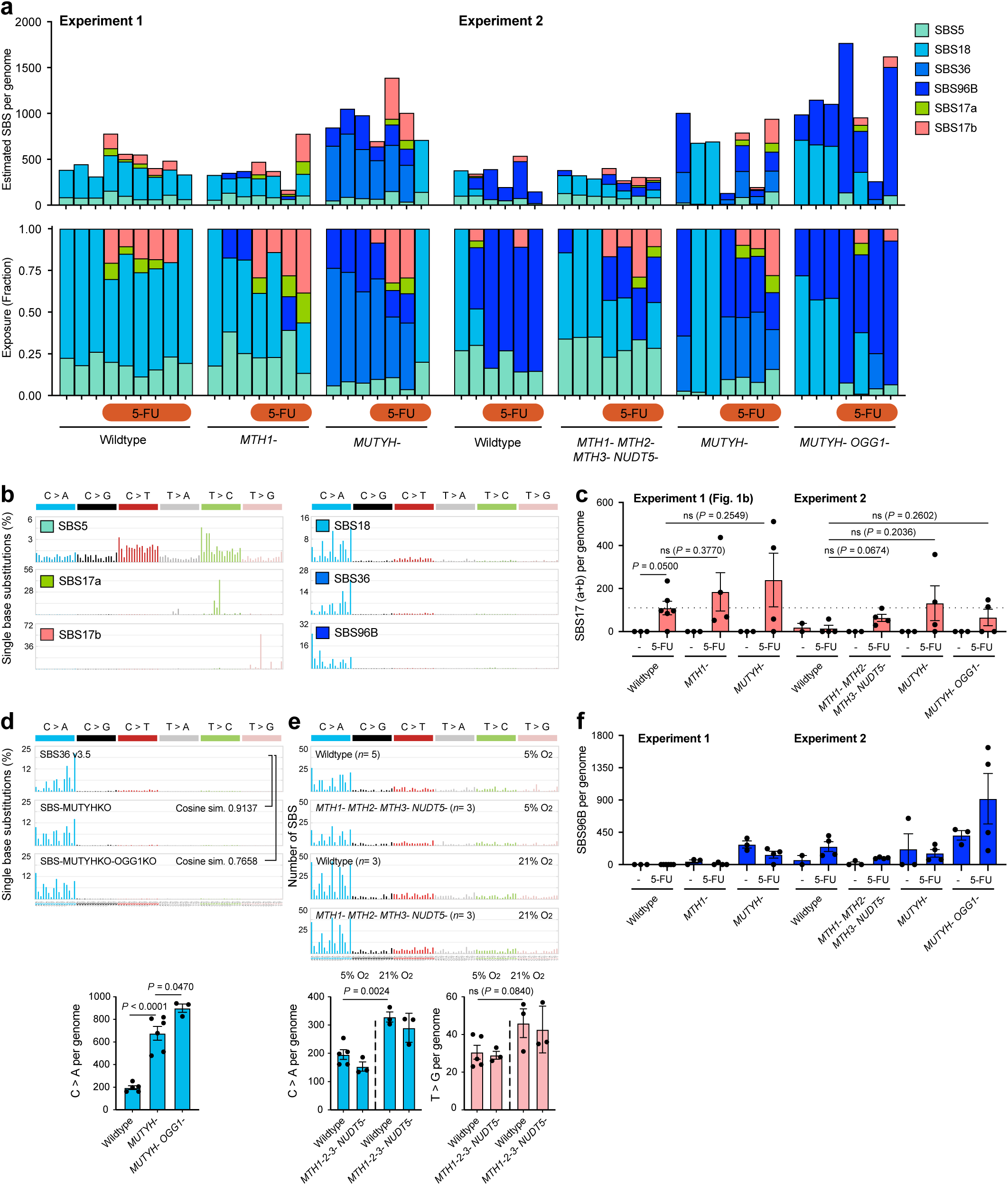
Mutation signature extraction in mutants linked to 8-oxo-guanine metabolism. **a)** Assignment of mutational signatures using SigProfiler toolkit. Stacked bar plots showing estimated number (top) and proportion (bottom) of each mutational signature in individual clones. **b)** Pattern of known COSMIC mutational signatures and the novel signature SBS96B, 96-classes of SBSs considering the six mutation types but also the bases immediately 5’ and 3’ of the mutated base. **c)** Burden of SBS17 mutations per (haploid) genome (*P* calculated by two-tailed unpaired *t* test, data shown as mean and s.e.m., *n* = 3-6). **d)** Top, comparison of mutational signatures caused by *MUTYH* deficiency found in cancer (SBS36, COSMIC v3.5) and our experimental signatures of *MUTYH* and *MUTYH/OGG1* deficiency. Bottom, burden of C>A mutations associated with the accumulation of oxidised guanine (*P* calculated by two-tailed unpaired *t* test, data shown as mean and s.e.m., *n* = 3-6). **e)** Mutational spectra and SBS burden in cells deficient in 4 guanine) but lack of T>G mutations associated with oxidation of the dGTP pool. (*P* calculated by two-tailed unpaired *t* test, data shown as mean and s.e.m., *n* = 3-5). **f)** Burden of a novel C>A signature (SBS96B) per (haploid) genome (*P* calculated by two-tailed unpaired *t* test, data shown as mean and s.e.m., *n* = 3-6).

**Extended Data Fig. 4.**
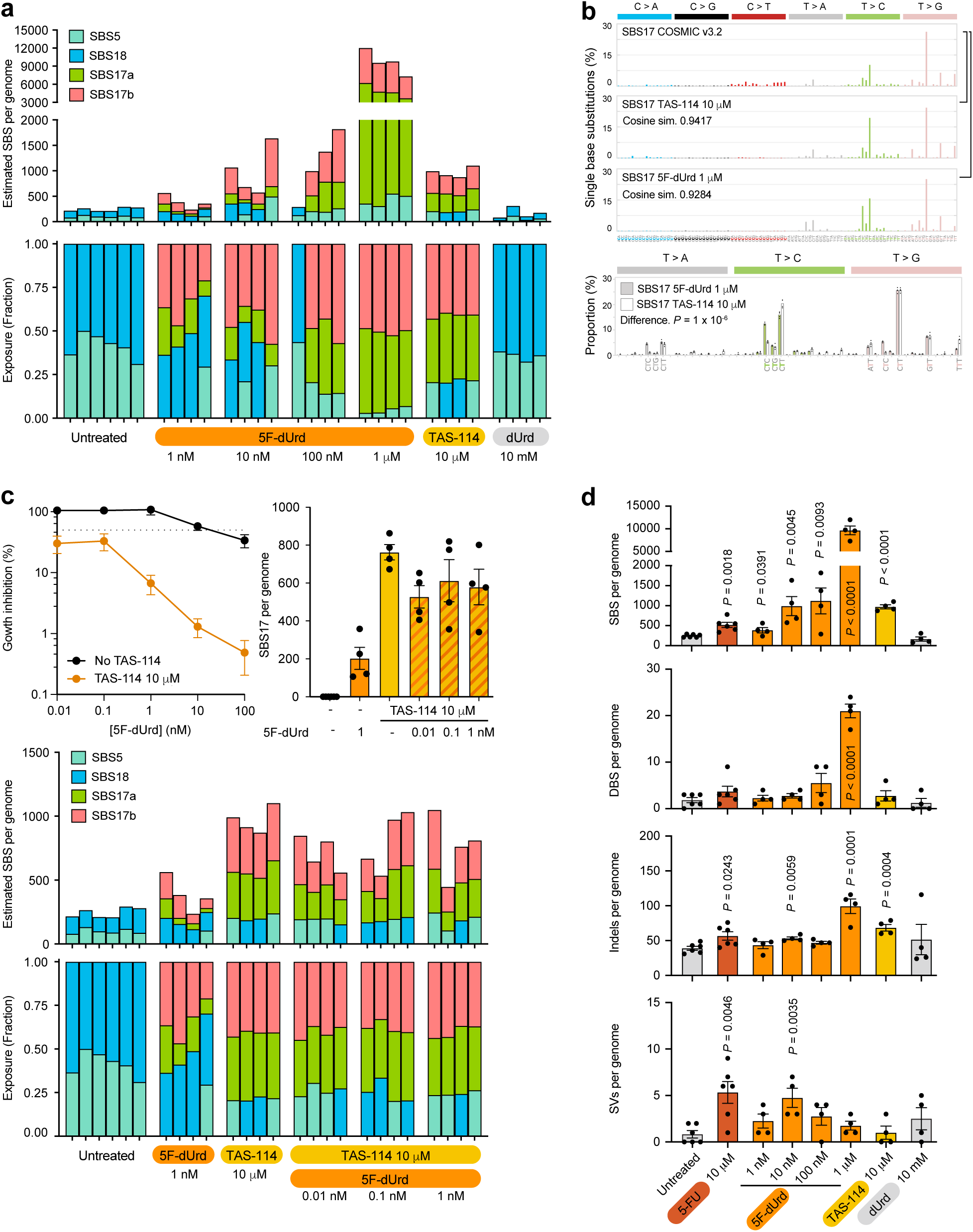
Mutation signatures caused by dUTP imbalance. **a)** Assignment of mutational signatures using SigProfiler toolkit. Stacked bar plots showing estimated number (top) and proportion (bottom) of each mutational signature in individual clones. **b)** Comparison of experimentally induced mutational signatures with SBS17 (COSMIC v3.2) found in cancer. Bottom, the trinucleotide context frequencies of T>N mutations induced by 5F-dUrd 1 M and TAS-114 10 M treatments, after subtracting mean background mutations (mean as bars, individual data points as dots, *n* = 4 per condition). *P* value from a parametric bootstrap test against a Dirichlet-multinomial null. **c)** Top left, growth inhibition assay showing the response of wildtype HAP1 cells to the 2′-deoxynucleoside 5F-dUridine (5F-dUrd) with or without the addition of the DUT inhibitor TAS-114. Average of three experiments each carried out in duplicate (data shown as mean and s.e.m., *n* = 3). Top right, burden of SBS17 mutations per (haploid) genome (*P* calculated by two-tailed unpaired *t* test, data shown as mean and s.e.m., *n* = 4-6). Bottom, assignment of mutational signatures using SigProfiler toolkit. Stacked bar plots showing estimated number (top) and proportion (bottom) of each mutational signature in individual clones. **d)** Burden of single-base substitutions (SBS), doublet-base substitutions (DBS), insertions and deletions, and structural variants (SVs) per (haploid) genome (*P* calculated by two-tailed unpaired *t* test, data shown as mean and s.e.m., *n* = 4-6).

**Extended Data Fig. 5.**
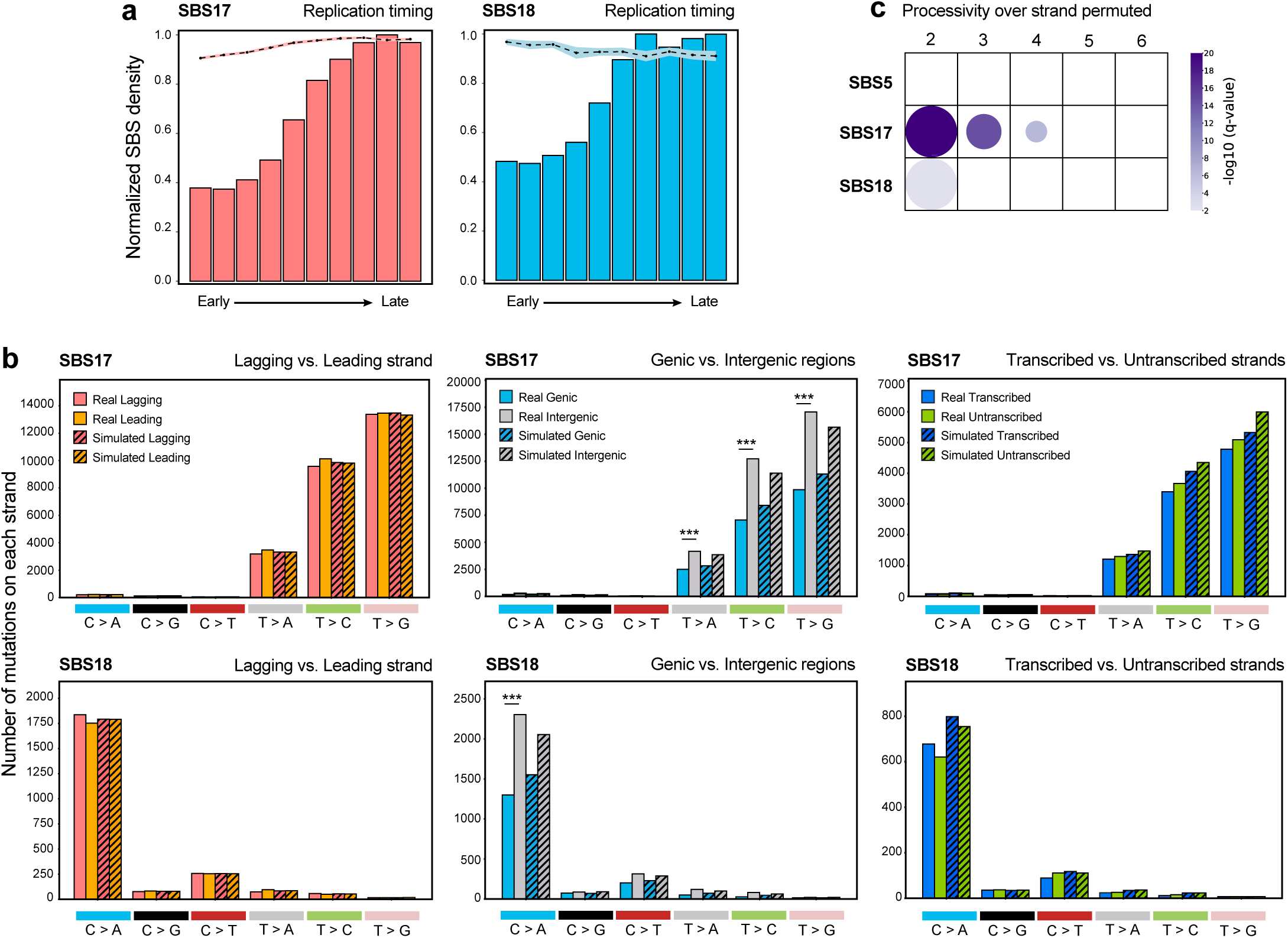
Topography of experimentally induced SBS17 mutations. **a)** Normalised mutational densities from early to late replicating regions in the mouse genome are shown with respect to real somatic mutations and simulated mutations. The line reflects the mean of simulated mutations, error bars represent the 95% confidence interval of this mean, the bars represent the number for real experimentally-induced SBS17 mutations or SBS18 mutations (HAP1 background) for comparison. **b)** Strand asymmetries SBS17 and SBS18 mutations. Bar plots display the number of mutations accumulated on each strand for six substitution subtypes based on the mutated pyrimidine base C>A, C>G, C>T, T>A, T>C, and T>G. Simulated mutations on the same strands are displayed in shaded bar plots. Statistically significant strand asymmetries are shown (*P* calculated by Fisher’s exact test corrected for multiple testing using Benjamini-Hochberg). **c)** Strand-coordinated mutagenesis of mutational signatures. Dot size proportional to number of strand-coordinated mutation clusters per cluster size, colour indicates –log10(q-value); q-value represents the Benjamini-Hochberg corrected p-value from a one-sided z-test against the strand-permuted null.

**Extended Data Fig. 6.**
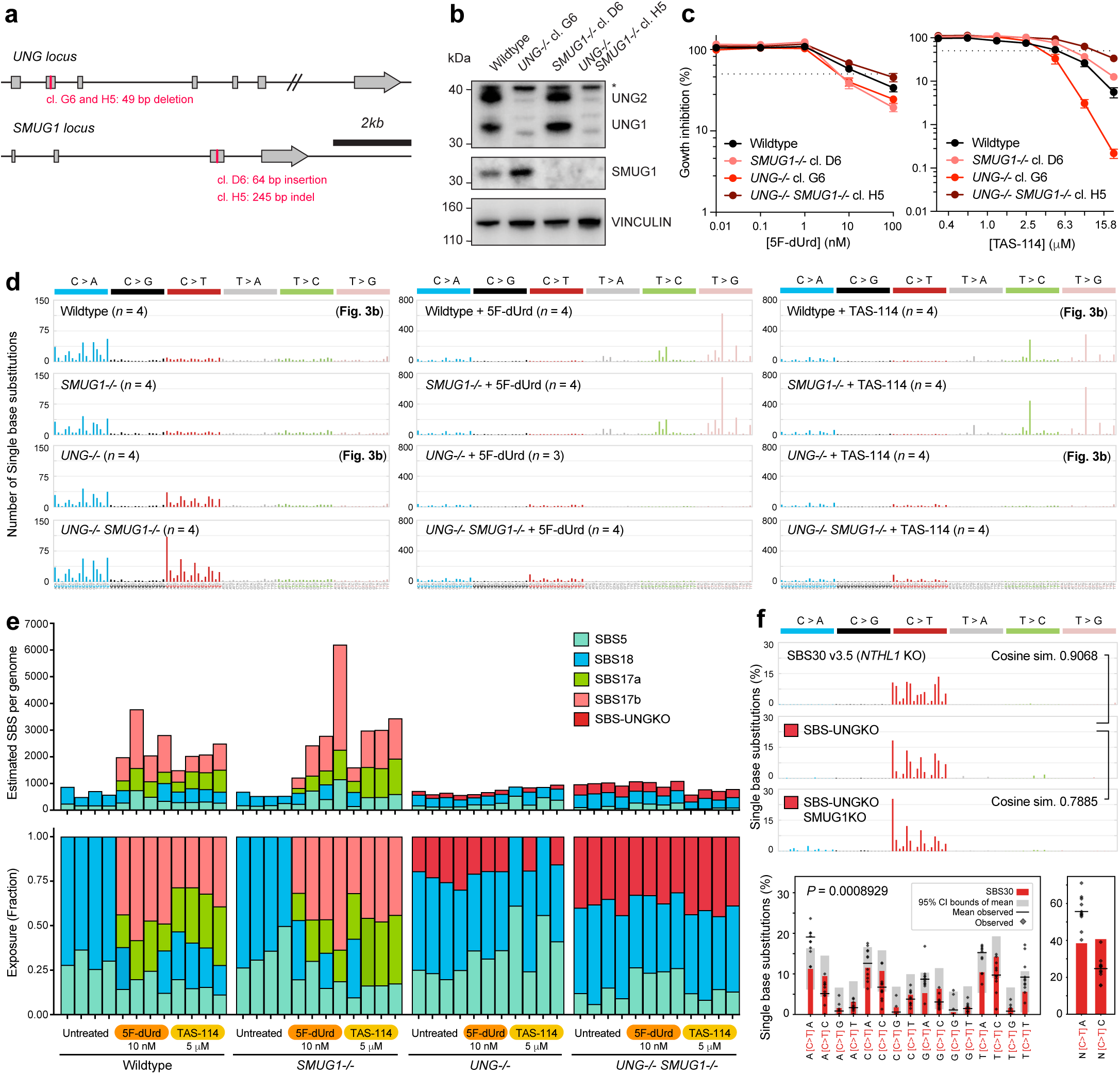
Generation of uracil glycosylase-deficient HAP1 mutants and signature extraction. **a)** Schematic cartoon of the human *loci*, targeted exons and introduced genetic modifications. See **Table 1** for sequence information and additional details. **b)** Validation of putative knockout lines by Western blot. **c)** Growth inhibition assay showing the response of HAP1 mutants to the 2′-deoxynucleoside 5F-dUridine (5F-dUrd) or the DUT inhibitor TAS-114. Average of three experiments each carried out in duplicate (data shown as mean and s.e.m., *n* = 3). **d)** Full dataset 96-classes of single-base substitutions (SBSs) considering the six mutation types but also the bases immediately 5’ and 3’ of the mutated base. Each graph represents the average mutation pattern for *n* genomes, where *n* is indicated in each panel. Panels also shown in Fig. 3b are indicated. **e)** Assignment of mutational signatures using SigProfiler toolkit. Stacked bar plots showing estimated number (top) and proportion (bottom) of each mutational signature in individual clones. **f)** Top, comparison of trinucleotide context of signatures caused by *NTHL1* deficiency found in cancer (SBS30, COSMIC v3.5) and our experimental signatures of *UNG* and *UNG/SMUG1* deficiency, after subtracting mean wildtype background mutations. Bottom, C>T component of UNG deficiency and SBS30 signatures (left) and nCa and nCc frequencies (right). Family-wise error rate -corrected *P*-value and 95% CI from a parametric bootstrap test against a Dirichlet-multinomial null (*n* = 11).

**Extended Data Fig. 7.**
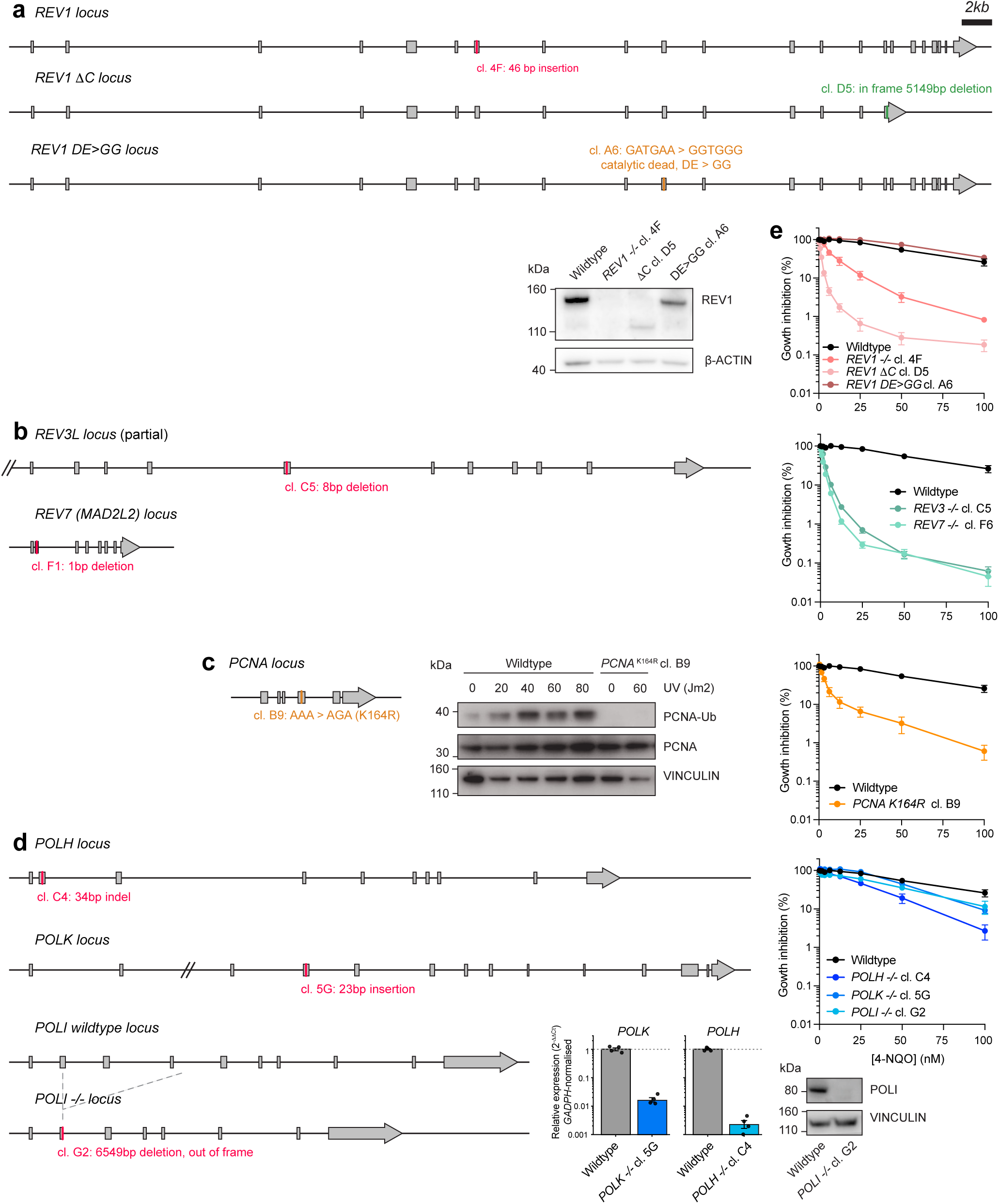
Generation and validation of mutants deficient in translesion synthesis (TLS). **a)** Schematic cartoon of the human *REV1 locus*, targeted exons and introduced genetic modifications. Validation of putative knockout and mutant lines by Western blot. **b)** Schematic cartoon of the human *REV3* and *REV7 loci*, which code for subunits of Pol. Validation in **e)** by sensitivity to 4-NQO. **c)** Schematic cartoon of the human *PCNA locus*, introduction of a point mutation in the endogenous locus by base editing. Validation by Western blot, showing lack of PCNA monoubiquitination in response to UV irradiation. **d)** Schematic cartoon of the human *POLH, POLK* and *POLI loci*. Validation by quantitative PCR of mRNA or Western blot. **a-d)** See **Table 1** for sequence information and additional details. **e)** Growth inhibition assay showing the response of TLS-deficient HAP1 cells to 4-nitroquinoline 1-oxide) (4-NQO), which creates bulky lesions. Average of three experiments carried out in duplicate (data shown as mean and s.e.m., *n* = 3).

**Extended Data Fig. 8.**
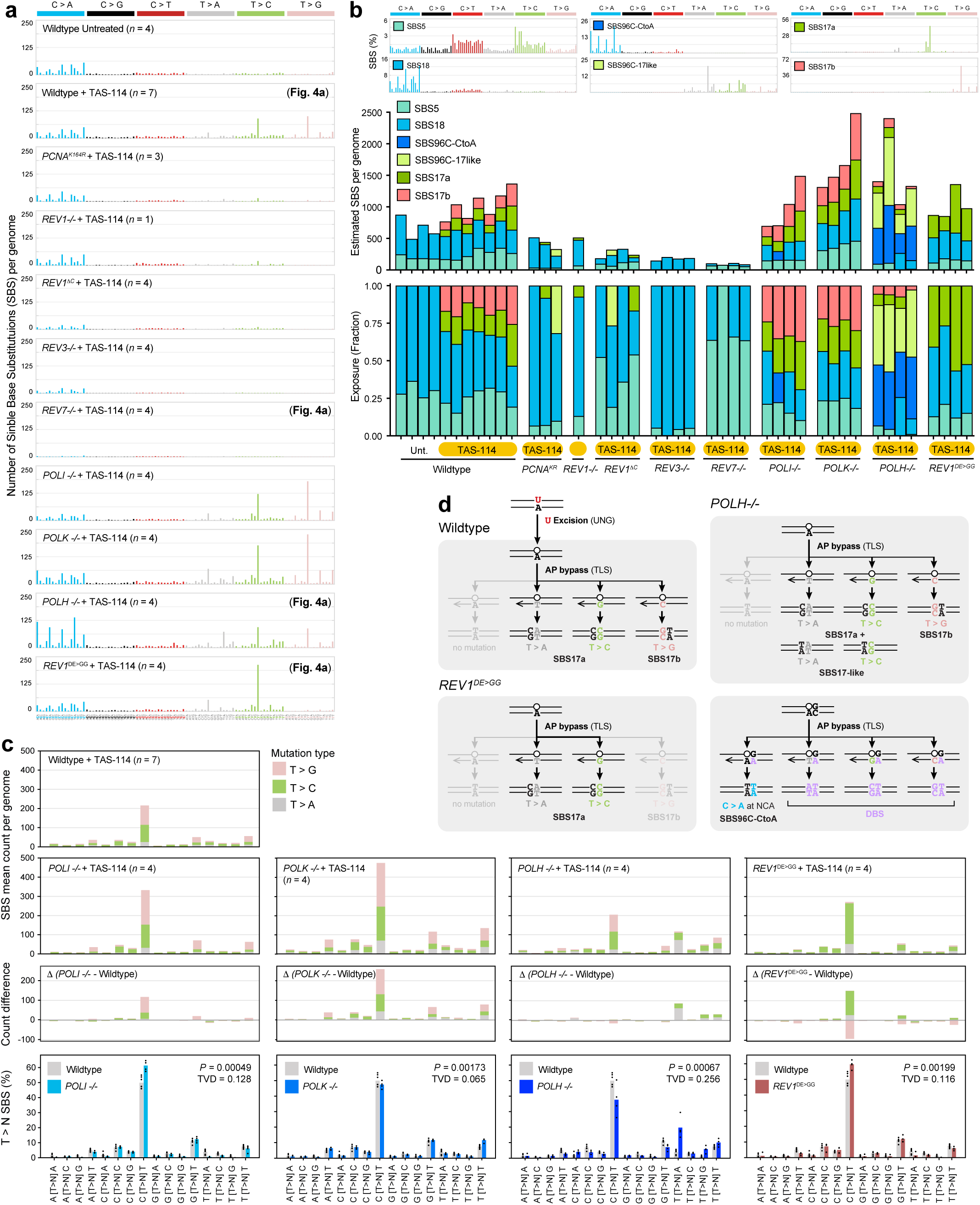
Extraction of substitutions signatures in mutants deficient in translesion synthesis (TLS). **a)** Full dataset 96-classes of single-base substitutions (SBSs) considering the six mutation types but also the bases immediately 5’ and 3’ of the mutated base. Each graph represents the average mutation pattern for *n* genomes, where *n* is indicated in each panel. Panels also shown in Fig. 4a are indicated. **b)** Assignment of mutational signatures using SigProfiler toolkit. Stacked bar plots showing estimated number (top) and proportion (bottom) of each mutational signature in individual clones. **c)** T>N mutation counts across trinucleotide contexts for different TAS-114 treated genotypes. Δ-panels display counts after subtracting wildtype from each genotype. Bottom, T>N trinucleotide mutation frequencies induced by TAS-114, comparing mutant phenotypes to wildtype. The total variation distance (TVD) is a measure of difference between two probability distributions, ranging from 0 (identical) to 1 (completely disjoint). Mean as bars, individual data points as dots, *n* = 4 per genotype, *P*-values from parametric bootstrap tests against a Dirichlet-multinomial null; *n*=7 for wildtype, *n*=4 per mutant genotype. **d)** Schematic cartoon summarising our results for the generation of single and doublet-base substitutions (DBS) and associated mutational signatures, in wildtype and TLS-deficient mutants.

**Extended Data Fig. 9.**
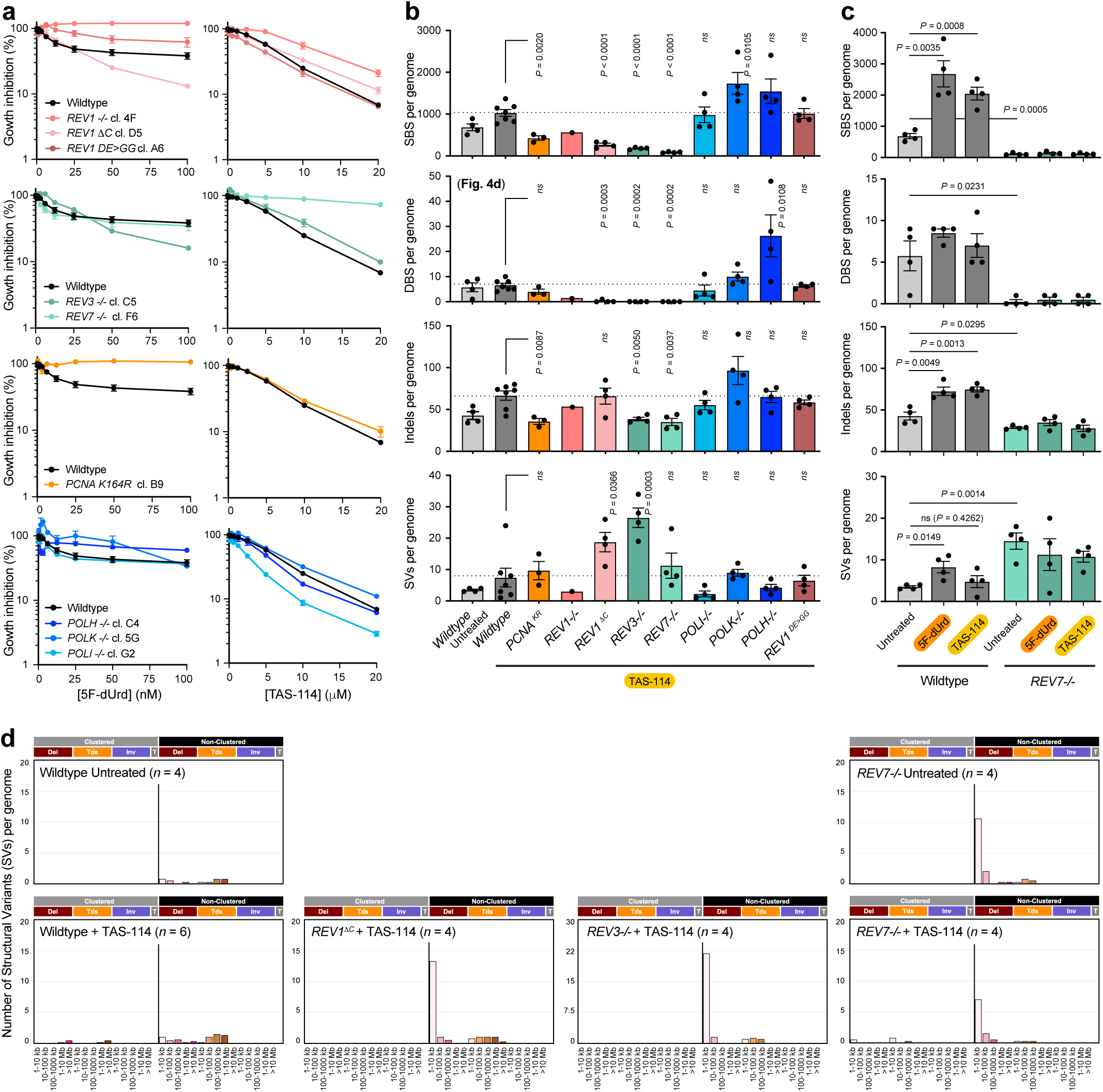
Structural variants in mutants deficient in translesion synthesis (TLS). **a)** Growth inhibition assay showing the response of TLS-deficient HAP1 cells to the 2′-deoxynucleoside 5F-dUridine (5F-dUrd) or the DUT inhibitor TAS-114. Average of two experiments carried out in duplicate (data shown as mean and s.e.m., *n* = 2). **b)** Burden of single-base substitutions (SBS), doublet-base substitutions (DBS), insertions and deletions and structural variants (SV) per (diploid) genome, for a panel of TLS-deficient mutants treated with TAS-114 10 M (*P* calculated by two-tailed unpaired *t* test, data shown as mean and s.e.m., *n* = 3-6, but *n* = 1 for *REV1-/-*). **c)** Burden of single-base substitutions (SBS), doublet-base substitutions (DBS), insertions and deletions and structural variants (SV) per (diploid) genome, for a panel of wildtype or *REV7-/-* treated with 5F-dUrd 10 nM or TAS-1145 M (*P* calculated by two-tailed unpaired *t* test, data shown as mean and s.e.m., *n* = *n* = 3-6, but *n* = 1 for *REV1-/-*). **d)** Pattern of large structural variants (SVs), following the classification from the COSMIC database. Each graph represents the average mutation pattern for *n* genomes, where *n* is indicated in each panel.

**Extended Data Fig. 10.**
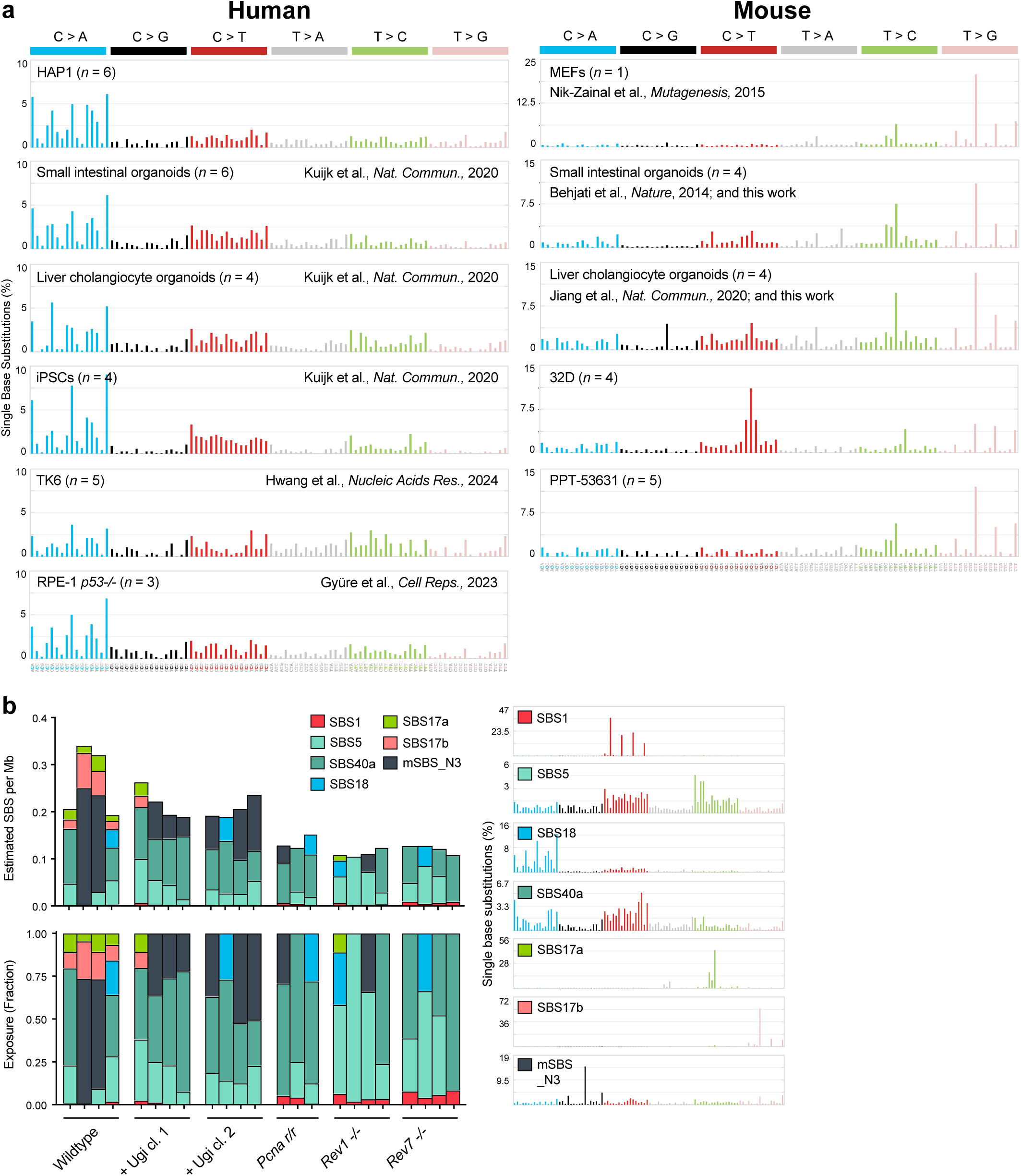
Mouse cells accumulate SBS17 mutations spontaneously *in vitro* which share the same mechanism. **a)** Comparison of mutational spectra across human and mouse cells grown *in vitro*, either previously published or generated for this study. 96-classes of single-base substitutions (SBSs) considering the six mutation types but also the bases immediately 5’ and 3’ of the mutated base. Each graph represents the average mutation pattern for *n* genomes, where *n* is indicated in each panel. **b)** Assignment of mutational signatures using SigProfiler toolkit. Stacked bar plots showing estimated number (top) and proportion (bottom) of each mutational signature in individual subcultures. Right, pattern of known COSMIC mutational signatures and the mouse-specific signature mSBS_N3 published previously (Riva et al *Nat. Genetics* 2020, ref ^69^), 96-classes of SBSs considering the six mutation types but also the bases immediately 5’ and 3’ of the mutated base.

## References

1. Blokzijl, F. et al. Tissue-specific mutation accumulation in human adult stem cells during life. Nature 538, 260–264 (2016).

2. Martincorena, I. & Campbell, P. J. Somatic mutation in cancer and normal cells. Science 349, 1483–9 (2015).

3. Alexandrov, L. B. et al. Signatures of mutational processes in human cancer. Nature 500, 415–421 (2013).

4. Alexandrov, L. B. et al. The repertoire of mutational signatures in human cancer. Nature 578, 94–101 (2020).

5. Dulak, A. M. et al. Exome and whole-genome sequencing of esophageal adenocarcinoma identifies recurrent driver events and mutational complexity. Nat. Genet. 45, 478–486 (2013).

6. Secrier, M. et al. Mutational signatures in esophageal adenocarcinoma define etiologically distinct subgroups with therapeutic relevance. Nat. Genet. 48, 1131–1141 (2016).

7. Busslinger, G. A. et al. Molecular characterization of Barrett’s esophagus at single-cell resolution. Proc. Natl. Acad. Sci. U. S. A. 118, (2021).

8. Abbas, S. et al. Mutational signature dynamics shaping the evolution of oesophageal adenocarcinoma. Nat. Commun. 14, 4239 (2023).

9. Moore, L. et al. The mutational landscape of human somatic and germline cells. Nature 10.1038/s41586-021-03822-7 (2021) doi:10.1038/s41586-021-03822-7.

10. Poetsch, A. R. The genomics of oxidative DNA damage, repair, and resulting mutagenesis. Comput. Struct. Biotechnol. J. 18, 207–219 (2020).

11. van der Ham, C. G. et al. Mutational mechanisms in multiply relapsed pediatric acute lymphoblastic leukemia. Leukemia 38, 2366–2375 (2024).

12. Tomkova, M., Tomek, J., Kriaucionis, S. & Schuster-Böckler, B. Mutational signature distribution varies with DNA replication timing and strand asymmetry. Genome Biol. 19, 129 (2018).

13. Dvorak, K. et al. Bile acids in combination with low pH induce oxidative stress and oxidative DNA damage: relevance to the pathogenesis of Barrett’s oesophagus. Gut 56, 763–71 (2007).

14. Foster, P. L., Lee, H., Popodi, E., Townes, J. P. & Tang, H. Determinants of spontaneous mutation in the bacterium Escherichia coli as revealed by whole-genome sequencing. Proc. Natl. Acad. Sci. U. S. A. 112, E5990–9 (2015).

15. Tajiri, T., Maki, H. & Sekiguchi, M. Functional cooperation of MutT, MutM and MutY proteins in preventing mutations caused by spontaneous oxidation of guanine nucleotide in Escherichia coli. Mutat. Res. 336, 257–67 (1995).

16. Longley, D. B., Harkin, D. P. & Johnston, P. G. 5-fluorouracil: mechanisms of action and clinical strategies. Nat. Rev. Cancer 3, 330–8 (2003).

17. Vodenkova, S. et al. 5-fluorouracil and other fluoropyrimidines in colorectal cancer: Past, present and future. Pharmacol. Ther. 206, 107447 (2020).

18. Christensen, S. et al. 5-Fluorouracil treatment induces characteristic T to G mutations in human cancer. Nat. Commun. 10, 4571 (2019).

19. Pich, O. et al. The mutational footprints of cancer therapies. Nat. Genet. 51, 1732–1740 (2019).

20. Focaccetti, C. et al. Effects of 5-fluorouracil on morphology, cell cycle, proliferation, apoptosis, autophagy and ROS production in endothelial cells and cardiomyocytes. PLoS One 10, e0115686 (2015).

21. Negrei, C. et al. Colon Cancer Cells Gene Expression Signature As Response to 5-Fluorouracil, Oxaliplatin, and Folinic Acid Treatment. Front. Pharmacol. 7, 172 (2016).

22. Carreras-Puigvert, J. et al. A comprehensive structural, biochemical and biological profiling of the human NUDIX hydrolase family. Nat. Commun. 8, 1541 (2017).

23. Hori, M., Satou, K., Harashima, H. & Kamiya, H. Suppression of mutagenesis by 8-hydroxy-2’-deoxyguanosine 5’-triphosphate (7,8-dihydro-8-oxo-2’-deoxyguanosine 5’-triphosphate) by human MTH1, MTH2, and NUDT5. Free Radic. Biol. Med. 48, 1197–201 (2010).

24. Ishibashi, T., Hayakawa, H. & Sekiguchi, M. A novel mechanism for preventing mutations caused by oxidation of guanine nucleotides. EMBO Rep. 4, 479–83 (2003).

25. Takagi, Y. et al. Human MTH3 (NUDT18) protein hydrolyzes oxidized forms of guanosine and deoxyguanosine diphosphates: comparison with MTH1 and MTH2. J. Biol. Chem. 287, 21541–9 (2012).

26. Hashiguchi, K., Hayashi, M., Sekiguchi, M. & Umezu, K. The roles of human MTH1, MTH2 and MTH3 proteins in maintaining genome stability under oxidative stress. Mutat. Res. 808, 10–19 (2018).

27. Viel, A. et al. A Specific Mutational Signature Associated with DNA 8-Oxoguanine Persistence in MUTYH-defective Colorectal Cancer. EBioMedicine 20, 39–49 (2017).

28. Pilati, C. et al. Mutational signature analysis identifies MUTYH deficiency in colorectal cancers and adrenocortical carcinomas. Journal of Pathology 242, 10–15 (2017).

29. Chen, J.-K. et al. An RNA Damage Response Network Mediates the Lethality of 5-FU in Clinically Relevant Tumor Types. Preprint at 10.1101/2023.04.28.538590 (2023).

30. Yano, W. et al. TAS-114, a First-in-Class Dual dUTPase/DPD Inhibitor, Demonstrates Potential to Improve Therapeutic Efficacy of Fluoropyrimidine-Based Chemotherapy. Mol. Cancer Ther. 17, 1683–1693 (2018).

31. el-Hajj, H. H., Zhang, H. & Weiss, B. Lethality of a dut (deoxyuridine triphosphatase) mutation in Escherichia coli. J. Bacteriol. 170, 1069–75 (1988).

32. Gadsden, M. H., McIntosh, E. M., Game, J. C., Wilson, P. J. & Haynes, R. H. dUTP pyrophosphatase is an essential enzyme in Saccharomyces cerevisiae. EMBO J. 12, 4425–31 (1993).

33. Otlu, B. et al. Topography of mutational signatures in human cancer. Cell Rep. 42, (2023).

34. Arnedo-Pac, C., Muiños, F., Gonzalez-Perez, A. & Lopez-Bigas, N. Hotspot propensity across mutational processes. Mol. Syst. Biol. 20, 6–27 (2024).

35. Roberts, S. A. et al. Clustered mutations in yeast and in human cancers can arise from damaged long single-strand DNA regions. Mol. Cell 46, 424–35 (2012).

36. Takahashi, I. & Marmur, J. Replacement of Thymidylic Acid by Deoxyuridylic Acid in the Deoxyribonucleic Acid of a Transducing Phage for Bacillus subtilis. Nature 197, 794–795 (1963).

37. Lindahl, T. Instability and decay of the primary structure of DNA. Nature 362, 709–15 (1993).

38. Lindahl, T. & Nyberg, B. Heat-induced deamination of cytosine residues in deoxyribonucleic acid. Biochemistry 13, 3405–10 (1974).

39. Lindahl, T. An N-glycosidase from Escherichia coli that releases free uracil from DNA containing deaminated cytosine residues. Proc. Natl. Acad. Sci. U. S. A. 71, 3649–53 (1974).

40. Krokan, H. E. & Bjørås, M. Base excision repair. Cold Spring Harb. Perspect. Biol. 5, a012583 (2013).

41. Musiani, D. et al. Uracil processing by SMUG1 in the absence of UNG triggers homologous recombination and selectively kills BRCA1/2-deficient tumors. Mol. Cell 85, 1072–1084.e10 (2025).

42. Dingler, F. A., Kemmerich, K., Neuberger, M. S. & Rada, C. Uracil excision by endogenous SMUG1 glycosylase promotes efficient Ig class switching and impacts on A:T substitutions during somatic mutation. Eur. J. Immunol. 44, 1925–35 (2014).

43. Kemmerich, K., Dingler, F. A., Rada, C. & Neuberger, M. S. Germline ablation of SMUG1 DNA glycosylase causes loss of 5-hydroxymethyluracil- and UNG-backup uracil-excision activities and increases cancer predisposition of Ung-/-Msh2-/-mice. Nucleic Acids Res. 40, 6016–25 (2012).

44. Grolleman, J. E. et al. Mutational Signature Analysis Reveals NTHL1 Deficiency to Cause a Multi-tumor Phenotype. Cancer Cell 35, 256–266.e5 (2019).

45. Drost, J. et al. Use of CRISPR-modified human stem cell organoids to study the origin of mutational signatures in cancer. Science 358, 234–238 (2017).

46. Sale, J. E. Translesion DNA synthesis and mutagenesis in eukaryotes. Cold Spring Harb. Perspect. Biol. 5, a012708 (2013).

47. Ross, A.-L., Simpson, L. J. & Sale, J. E. Vertebrate DNA damage tolerance requires the C-terminus but not BRCT or transferase domains of REV1. Nucleic Acids Res. 33, 1280–9 (2005).

48. Nelson, J. R., Lawrence, C. W. & Hinkle, D. C. Deoxycytidyl transferase activity of yeast REV1 protein. Nature 382, 729–731 (1996).

49. Gyüre, Z. et al. Spontaneous mutagenesis in human cells is controlled by REV1-Polymerase ζ and PRIMPOL. Cell Rep. 42, 112887 (2023).

50. Abe, T., Branzei, D. & Hirota, K. DNA Damage Tolerance Mechanisms Revealed from the Analysis of Immunoglobulin V Gene Diversification in Avian DT40 Cells. Genes (Basel). 9, (2018).

51. Zhang, H. & Lawrence, C. W. The error-free component of the RAD6/RAD18 DNA damage tolerance pathway of budding yeast employs sister-strand recombination. Proc. Natl. Acad. Sci. U. S. A. 102, 15954–9 (2005).

52. Jiang, Y. et al. Tissue-specific mutagenesis from endogenous guanine damage is suppressed by Polκ and DNA repair. Nat. Commun. 17, 436 (2025).

53. Nik-Zainal, S. et al. The genome as a record of environmental exposure. Mutagenesis 30, 763–770 (2015).

54. Behjati, S. et al. Genome sequencing of normal cells reveals developmental lineages and mutational processes. Nature 513, 422–425 (2014).

55. Hwang, T. et al. Comprehensive whole-genome sequencing reveals origins of mutational signatures associated with aging, mismatch repair deficiency and temozolomide chemotherapy. Nucleic Acids Res. 53, (2025).

56. Kuijk, E. et al. The mutational impact of culturing human pluripotent and adult stem cells. Nat. Commun. 11, 2493 (2020).

57. Petljak, M. et al. Characterizing Mutational Signatures in Human Cancer Cell Lines Reveals Episodic APOBEC Mutagenesis. Cell 176, 1282–1294.e20 (2019).

58. Di Noia, J. & Neuberger, M. S. Altering the pathway of immunoglobulin hypermutation by inhibiting uracil-DNA glycosylase. Nature 419, 43–8 (2002).

59. Rada, C. et al. Immunoglobulin isotype switching is inhibited and somatic hypermutation perturbed in UNG-deficient mice. Curr. Biol. 12, 1748–55 (2002).

60. Simpson, L. J. & Sale, J. E. Rev1 is essential for DNA damage tolerance and non-templated immunoglobulin gene mutation in a vertebrate cell line. EMBO J. 22, 1654–64 (2003).

61. Nik-Zainal, S. et al. Mutational processes molding the genomes of 21 breast cancers. Cell 149, 979–93 (2012).

62. Taylor, B. J. et al. DNA deaminases induce break-associated mutation showers with implication of APOBEC3B and 3A in breast cancer kataegis. Elife 2, e00534 (2013).

63. Petljak, M. et al. Mechanisms of APOBEC3 mutagenesis in human cancer cells. Nature 607, 799–807 (2022).

64. Machado, H. E. et al. Diverse mutational landscapes in human lymphocytes. Nature 608, 724–732 (2022).

65. Roerink, S. F. et al. Intra-tumour diversification in colorectal cancer at the single-cell level. Nature 556, 437–462 (2018).

66. Shapiro, R. & Klein, R. S. The deamination of cytidine and cytosine by acidic buffer solutions. Mutagenic implications. Biochemistry 5, 2358–62 (1966).

67. Doi, T. et al. First-in-human phase 1 study of novel dUTPase inhibitor TAS-114 in combination with S-1 in Japanese patients with advanced solid tumors. Invest. New Drugs 37, 507–518 (2019).

68. Kawazoe, A. et al. A multicenter phase II study of TAS-114 in combination with S-1 in patients with pretreated advanced gastric cancer (EPOC1604). Gastric Cancer 24, 190–196 (2021).

69. Riva, L. et al. The mutational signature profile of known and suspected human carcinogens in mice. Nat. Genet. 10.1038/s41588-020-0692-4 (2020) doi:10.1038/s41588-020-0692-4.

70. Cancer Genome Atlas Network. Comprehensive molecular characterization of human colon and rectal cancer. Nature 487, 330–7 (2012).

71. Cancer Genome Atlas Research Network. Comprehensive molecular characterization of gastric adenocarcinoma. Nature 513, 202–9 (2014).

72. Cancer Genome Atlas Research Network et al. Integrated genomic characterization of oesophageal carcinoma. Nature 541, 169–175 (2017).

73. Perdomo, S. et al. The Mutographs biorepository: A unique genomic resource to study cancer around the world. Cell genomics 4, 100500 (2024).

74. Martínez-Jiménez, F. et al. Pan-cancer whole-genome comparison of primary and metastatic solid tumours. Nature 618, 333–341 (2023).

75. Paulson, T. G. et al. Somatic whole genome dynamics of precancer in Barrett’s esophagus reveals features associated with disease progression. Nat. Commun. 13, 2300 (2022).

76. Díaz-Gay, M. et al. Assigning mutational signatures to individual samples and individual somatic mutations with SigProfilerAssignment. Bioinformatics 39, (2023).

77. Martínez-Jiménez, F. et al. A compendium of mutational cancer driver genes. Nat. Rev. Cancer 20, 555–572 (2020).

78. UniProt Consortium. UniProt: the Universal Protein Knowledgebase in 2023. Nucleic Acids Res. 51, D523–D531 (2023).

79. Geurts, M. H., et al. One-step generation of tumor models by base editor multiplexing in adult stem cell-derived organoids. Nat. Commun. 14, 4998 (2023).

80. Li, H. & Durbin, R. Fast and accurate short read alignment with Burrows-Wheeler transform. Bioinformatics 25, 1754–60 (2009).

81. Saunders, C. T. et al. Strelka: accurate somatic small-variant calling from sequenced tumor-normal sample pairs. Bioinformatics 28, 1811–7 (2012).

82. Cibulskis, K. et al. Sensitive detection of somatic point mutations in impure and heterogeneous cancer samples. Nat. Biotechnol. 31, 213–9 (2013).

83. Wardell, C. P., Ashby, C. & Bauer, M. A. FiNGS: high quality somatic mutations using filters for next generation sequencing. BMC Bioinformatics 22, 77 (2021).

84. Chen, X., et al. Manta: rapid detection of structural variants and indels for germline and cancer sequencing applications. Bioinformatics 32, 1220–2 (2016).

85. Cameron, D. L. et al. GRIDSS2: comprehensive characterisation of somatic structural variation using single breakend variants and structural variant phasing. Genome Biol. 22, 202 (2021).

86. Bergstrom, E. N. et al. SigProfilerMatrixGenerator: a tool for visualizing and exploring patterns of small mutational events. BMC Genomics 20, 685 (2019).

87. Islam, S. M. A. et al. Uncovering novel mutational signatures by de novo extraction with SigProfilerExtractor. Cell genomics 2, None (2022).

88. Otlu, B. & Alexandrov, L. B. Evaluating topography of mutational signatures with SigProfilerTopography. Genome Biol. 26, 134 (2025).

89. Klein, K. N. et al. Replication timing maintains the global epigenetic state in human cells. Science 372, 371–378 (2021).

90. Harrison, J. G., Calder, W. J., Shastry, V. & Buerkle, C. A. Dirichlet-multinomial modelling outperforms alternatives for analysis of microbiome and other ecological count data. Mol. Ecol. Resour. 20, 481–497 (2020).

91. Tusher, V. G., Tibshirani, R. & Chu, G. Significance analysis of microarrays applied to the ionizing radiation response. Proc. Natl. Acad. Sci. U. S. A. 98, 5116–21 (2001).

92. Westfall, P. H. & Young, S. S. Resampling-Based Multiple Testing: Examples and Methods for p-Value Adjustment. (Wiley-Interscience, 1993).

93. Jansen, J. G. et al. Strand-biased defect in C/G transversions in hypermutating immunoglobulin genes in Rev1-deficient mice. J. Exp. Med. 203, 319–23 (2006).

94. Langerak, P., Nygren, A. O. H., Krijger, P. H. L., van den Berk, P. C. M. & Jacobs, H. A/T mutagenesis in hypermutated immunoglobulin genes strongly depends on PCNAK164 modification. J. Exp. Med. 204, 1989–98 (2007).

95. Ghezraoui, H. et al. 53BP1 cooperation with the REV7-shieldin complex underpins DNA structure-specific NHEJ. Nature 560, 122–127 (2018).

96. Broutier, L. et al. Culture and establishment of self-renewing human and mouse adult liver and pancreas 3D organoids and their genetic manipulation. Nat. Protoc. 11, 1724–43 (2016).

97. Hendriks, D. et al. Engineered human hepatocyte organoids enable CRISPR-based target discovery and drug screening for steatosis. Nat. Biotechnol. 41, 1567–1581 (2023).

98. Fujii, M., Matano, M., Nanki, K. & Sato, T. Efficient genetic engineering of human intestinal organoids using electroporation. Nat. Protoc. 10, 1474–85 (2015).

99. Abascal, F. et al. Somatic mutation landscapes at single-molecule resolution. Nature 593, 405–410 (2021).

100. Tischler, G. & Leonard, S. biobambam: tools for read pair collation based algorithms on BAM files. Source Code Biol. Med. 9, 13 (2014).

101. Sarno, A. et al. Uracil-DNA glycosylase UNG1 isoform variant supports class switch recombination and repairs nuclear genomic uracil. Nucleic Acids Res. 47, 4569–4585 (2019).

